# A simple *Agrobacterium*-mediated stable transformation technique for the hornwort model *Anthoceros agrestis*

**DOI:** 10.1101/2021.01.07.425778

**Authors:** Eftychios Frangedakis, Manuel Waller, Tomoaki Nishiyama, Hirokazu Tsukaya, Xia Xu, Yuling Yue, Michelle Tjahjadi, Andika Gunadi, Joyce Van Eck, Fay-Wei Li, Péter Szövényi, Keiko Sakakibara

## Abstract

We have developed a simple *Agrobacterium-mediated* method for the stable transformation of the hornwort *Anthoceros agrestis*, the fifth bryophyte species for which a genetic manipulation technique becomes available. High transformation efficiency was achieved by using thallus tissue grown under low-light conditions. We generated a total of 216 transgenic *A. agrestis* lines expressing the β-Glucuronidase (GUS), cyan, green, and yellow fluorescent proteins under the control of the CaMV 35S promoter and several endogenous promoters. Nuclear and plasma membrane localization with multiple color fluorescent proteins was also confirmed. The transformation technique described here should pave the way for detailed molecular and genetic studies of hornwort biology, providing much needed insight into the molecular mechanisms underlying symbiosis, carbon-concentrating mechanism, RNA editing, and land plant evolution in general.

## Introduction

The hornworts are one of the three lineages of bryophytes that diverged from liverworts and mosses about 460 million years ago (Morris et al., 2018; One Thousand Plant Transcriptomes Initiative, 2019; Li et al., 2020). While having only around 220 extant species (Söderström et al., 2016), hornworts are key to address diverse questions about land plant evolution and terrestrialization. Hornworts display a unique combination of features (Frangedakis et al., 2020) such as a sporophyte, that is produced by an indeterminate basal meristem, and bears stomata similar to mosses and vascular plants (Renzaglia et al., 2017). In addition, it is the only extant land plant lineage (together with a few *Selaginella* species (Liu et al., 2020)), that has a single (or just a few) algal-like chloroplast(s) per cell. The chloroplasts resemble those of algae in that they may contain pyrenoids, a carbon-concentrating structure that is shared with many algal lineages (Villarreal and Renner, 2012; Li et al., 2017). Hornwort plastids are also unique by exhibiting the highest RNA editing rates amongst land plants (Yoshinaga, 1996; Yoshinaga, 1997; Kugita, 2003; Small et al., 2019). Finally, hornworts are among the very few plant lineages that can establish symbiotic relationships with both endophytic cyanobacteria (Renzaglia et al., 2009) and various glomeromycotina and mucoromycotina fungal partners (mycorrhiza) (Desirò et al., 2013).

*Anthoceros agrestis* has been established as an experimental model system for hornworts and two isolates are currently available (Oxford and Bonn) (Szövényi et al., 2015). *A. agrestis*, like other bryophytes, has a haploid-dominant life cycle through which the haploid gametophyte phase alternates with the diploid sporophyte phase. The life cycle of *A. agrestis* starts with the germination of the haploid spores which develop into an irregularly shaped thallus (Fig. 1 A-C). Sexual reproduction occurs through fusion of the egg (produced in archegonia) and the motile sperm (produced in antheridia). The resulting embryo develops within the gametophyte and gives rise to the sporophyte (Fig. 1 D). *A. agrestis* can be easily grown under laboratory conditions and its haploid-dominant life cycle makes genetic analysis straightforward. The nuclear genome of *A. agrestis* was recently sequenced (Li et al., 2020) which is one of the smallest genomes amongst land plants. The species is monoicous, with male and female reproductive organs produced by the same individual. The sexual life cycle of *A. agrestis* can be completed under laboratory conditions within approximately 2-3 months (Szövényi et al., 2015). However, transformation of *A. agrestis* was not feasible until recently, posing a major obstacle to the analysis of gene function in hornworts, and more generally to land plant evo-devo studies.

**Figure 1:**
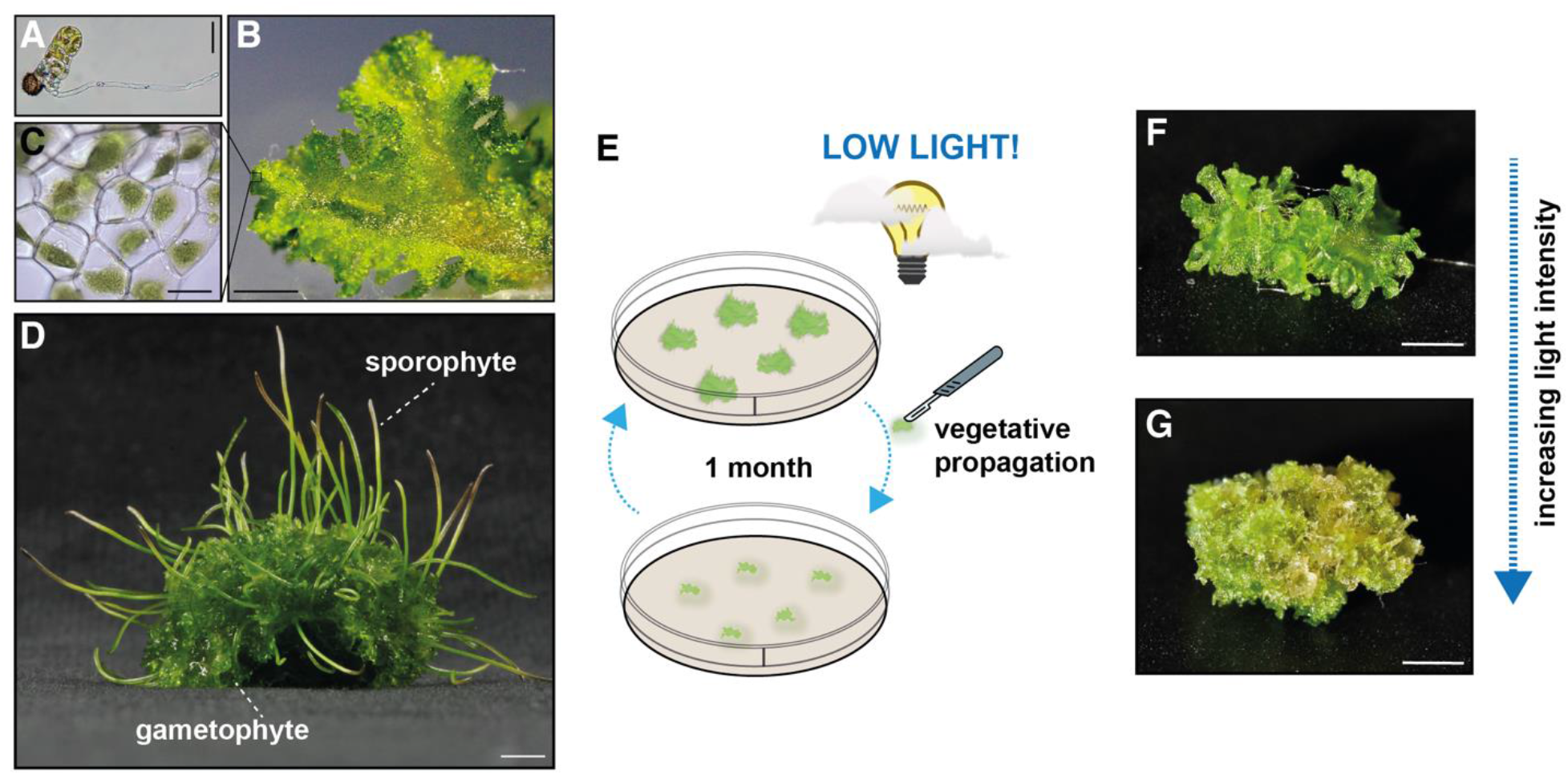
Morphological features of *Anthoceros agrestis* and effect of light on growth. A) Light micrograph (LM) of a germinating spore. Scale bar: 50 μm. B) Surface view of the irregularly shaped thallus (gametophyte). Scale bar: 1 mm. C) LM showing cells of mature gametophyte tissue with single chloroplasts. Scale bar: 10 μm. D) *A. agrestis* Oxford gametophyte with sporophytes. Scale bar: 4 mm. E) The conditions for the preparation of tissue used for transformation are critical. Plants must be propagated in axenic culture by transferring small thallus fragments (typically 1 x 1 mm) onto plates with fresh growth medium using sterile scalpels and then grown under low light conditions (3-5 μmol m^-2^ s^-1^) for 4 weeks. F) *A. agrestis* Oxford thallus tissue grown for 4 weeks under low light intensity (3-5 μmol m^-2^ s^-1^). G) *A. agrestis* Oxford thallus tissue grown for 4 weeks under high light intensity (80 μmol m^-2^ s^-1^). Scale bars: 1 mm.

Several approaches have been used for gene delivery in bryophytes, including polyethylene glycol (PEG)-mediated uptake of DNA by protoplasts (Schaefer et al., 1991), particle bombardment (Cho et al., 1999; Chiyoda et al., 2008) and *Agrobacterium tumefaciens-mediated* (revised scientific name *Rhizobium radiobacter* (Young et al., 2001)) transformation (Ishizaki et al., 2008; Kubota et al., 2013; Althoff and Zachgo, 2020). *Agrobacterium-mediated* transformation is a commonly used method for various plant species (Gelvin, 2003), it is relatively simple and does not require specialised or expensive equipment. In addition, *Agrobacterium-mediated* transformation has several advantages over other transformation methods, such as the integration of a lower number of transgene copies into the plant genome and the ability to transfer relatively large DNA segments with intact transgene genome integration.

In this study we report the first successful stable genetic transformation method for hornworts. The method is based on *Agrobacterium-mediated* transformation of *A. agrestis* thallus. We also report the successful expression and targeting of four different fluorescent proteins in two different cellular compartments, the plasma membrane and the nucleus. Finally, we characterize a number of native *A. agrestis* promoters for their potential to drive strong constitutive transgene expression that can be useful for future hornwort genetic studies.

## Results

We tested the potential of the *Agrobacterium-mediated* gene delivery method to recover stable *A. agrestis* transgenic lines. There are several critical factors that determine the efficiency of *Agrobacterium*-mediated gene delivery: (1) the selection of appropriate plant tissue for infection and conditions of tissue preparation, (2) the type and concentration of antibiotics applied to select for transgenic lines, (3) the choice of transformation vectors, (4) the choice of *Agrobacterium* strains, and (5) the conditions used for co-cultivation. Each of these factors is examined in this study.

### Selection of tissue and optimal growth conditions

*A. agrestis* gametophyte thallus was chosen as the appropriate tissue for transformation, because it is easily accessible, has a remarkable regenerative capacity and is haploid. *A. agrestis* thallus tissue cultures can easily be propagated and maintained by transfer of small thallus fragments (approximately 1 x 1 mm) onto fresh growth medium on a monthly basis (Supplemental Fig. S1). *A. agrestis* is similar to several other bryophytes in that an entire plant can regenerate from a small thallus fragment. This is in striking contrast with vascular plants, where usually the transition to an undifferentiated tissue state (callus), after treatment with extrinsic hormones such as auxin and cytokinin, is necessary before the regeneration of new plant tissue (Ikeuchi et al., 2013). The use of thallus has the additional advantage that the resulting transformants have a uniform genetic background.

We reasoned that similar to the liverwort *Marchantia polymorpha* (Kubota *et al*., 2013), fragmented tissue will be susceptible to *Agrobacterium* infection. In the case of *M. polymorpha*, the apical part of the thallus is removed to induce regeneration, followed by co-cultivation with *Agrobacterium* to generate transformed plants. *A. agrestis* thallus regeneration is similarly induced by fragmentation and presumably removal of the apical parts of the thallus. Consequently, a transformation approach similar to the one used for *M. polymorpha* was utilized and adapted to *A. agrestis*. However, unlike in *M. polymorpha*, the *A. agrestis* notch area and apical cells are not easily distinguishable, thus determining which part of the thallus should be removed is not easy. Therefore, we tested whether homogenization using dispensing tools is a suitable method for thallus tissue fragmentation. Different speed levels and duration of homogenization were examined. We found that homogenization of 0.5 g of thallus tissue in 15 mL of sterile water for 5 seconds (see Materials and methods) is sufficient to fragment the tissue without damaging the plants and results in rapid tissue regeneration.

In addition, we found that the light intensity used to cultivate *A. agrestis* tissue is a critical factor for successful transformation (Fig. 1 E-G). Tissue that was grown under low light conditions, even though smaller in size, had a more regular and flattened shape and was optimal for transformation. (Fig. 1 F-G and Supplemental Fig. S1 A). When tissue was grown under high light intensity (above 40 μmol m^-2^ s^-1^), no transformants were obtained.

### Selection of appropriate antibiotics

A selectable marker gene, most commonly one that confers antibiotic-resistance, is necessary for efficient recovery of stable transgenic lines following co-cultivation with *Agrobacterium*. To identify antibiotics and their appropriate concentration with cytotoxic effect on *A. agrestis*, untransformed *A. agrestis* (Oxford and Bonn) thallus fragments were subjected to different concentrations of hygromycin and geneticin/G418 (an analogue of neomycin and kanamycin). The tested concentrations ranged from 0 μg/mL - 20 μg/mL for hygromycin and 0 μg/mL - 150 μg/mL for geneticin/G418. We found that a 3-week incubation period with 10 μg/mL hygromycin was sufficient to inhibit growth of untransformed thallus tissue (Supplemental Fig. S2 and S3), whereas thallus tissue was not susceptible to geneticin/G418 even when supplied in high concentration (150 μg/mL) (Supplemental Fig. S4). Thus, we selected hygromycin as an appropriate selection agent for *A. agrestis* transformation.

### Preliminary tests with the GUS reporter

Preliminary transformation experiments were performed using the Oxford isolate and the pCAMBIA1305.2 plasmid containing the *hygromycin B phosphotransferase* gene (*hph*, conferring hygromycin resistance) driven by the Cauliflower mosaic virus 35S (CaMV 35S, hereafter called 35S) promoter, terminated with a 35S polyadenylation signal, and a *p-35S_s::GUSPlus* (β-Glucuronidase) transcription unit. *GUSPlus* contains a catalase intron to ensure that the observed GUS expression is not due to the *Agrobacterium*.

Homogenized regenerating thallus tissue was co-cultivated with the *Agrobacterium* AGL1 strain containing the pCAMBIA1305.2 plasmid, as well as *Agrobacterium* without a transformation vector as a negative control, in liquid KNOP media supplemented with sucrose. Media were also supplemented with 3’,5’-dimethoxy-4’-hydroxyacetophenone (acetosyringone) since phenolic compounds such as acetosyringone have been shown to be important for the virulence genes activation (Stachel et al., 1985). Co-cultivation duration was 3 days at 22°C on a shaker without any light supplementation (only ambient light from the room). After co-cultivation, the tissue was spread on solid KNOP plates supplemented with cefotaxime and hygromycin. After 3-4 weeks the tissue was transferred to fresh selective media (transformation outline in Fig. 2 and Supplemental Fig. S5). Emergence of rhizoids on surviving tissue fragments is a reliable indicator of successful transformation events (Fig. 3A-B). One to two months later successful transformants (plant fragments producing rhizoids) were visible. Finally, surviving plants were subjected to a third round of antibiotic selection to ensure false positives were eliminated. Thallus surviving selection on hygromycin exhibited GUS expression (Fig. 3C, Supplemental Fig. S6). No mock-transformed plants survived antibiotic selection; e.g. when transformation was carried out using *Agrobacterium* lacking the transformation vector. These results indicated that *A. agrestis* Oxford thallus tissue is susceptible to *Agrobacterium* infection and that the 35S promoter driving the *hph* gene is sufficient for selection of transformants.

**Figure 2:**
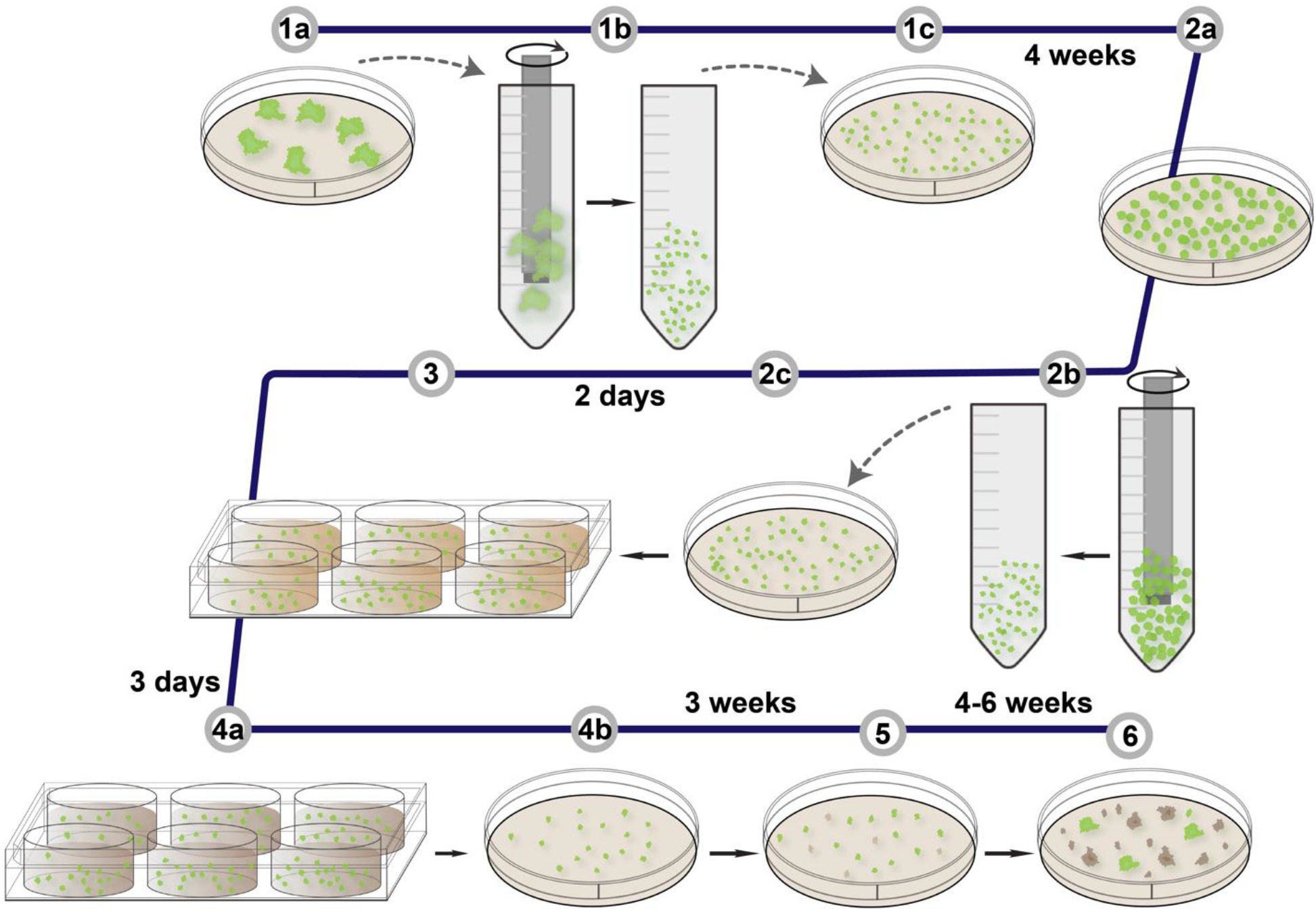
Outline of *Anthoceros agrestis* transformation method. **1a-c**: Tissue is homogenized, transferred on growth medium, and placed under low light conditions. **2a-c**: After 4 weeks, the tissue is homogenized again and grown for two additional days. **3**: The tissue is co-cultivated with *Agrobacterium* for three days and then **4a-b**: spread on appropriate antibiotic-containing growth medium. **5**: After 3 weeks, the tissue is transferred again onto freshly prepared antibiotic-containing growth medium for a second round of selection. **6**: After approximately 4-8 weeks, putative transformants are visible.

**Figure 3:**
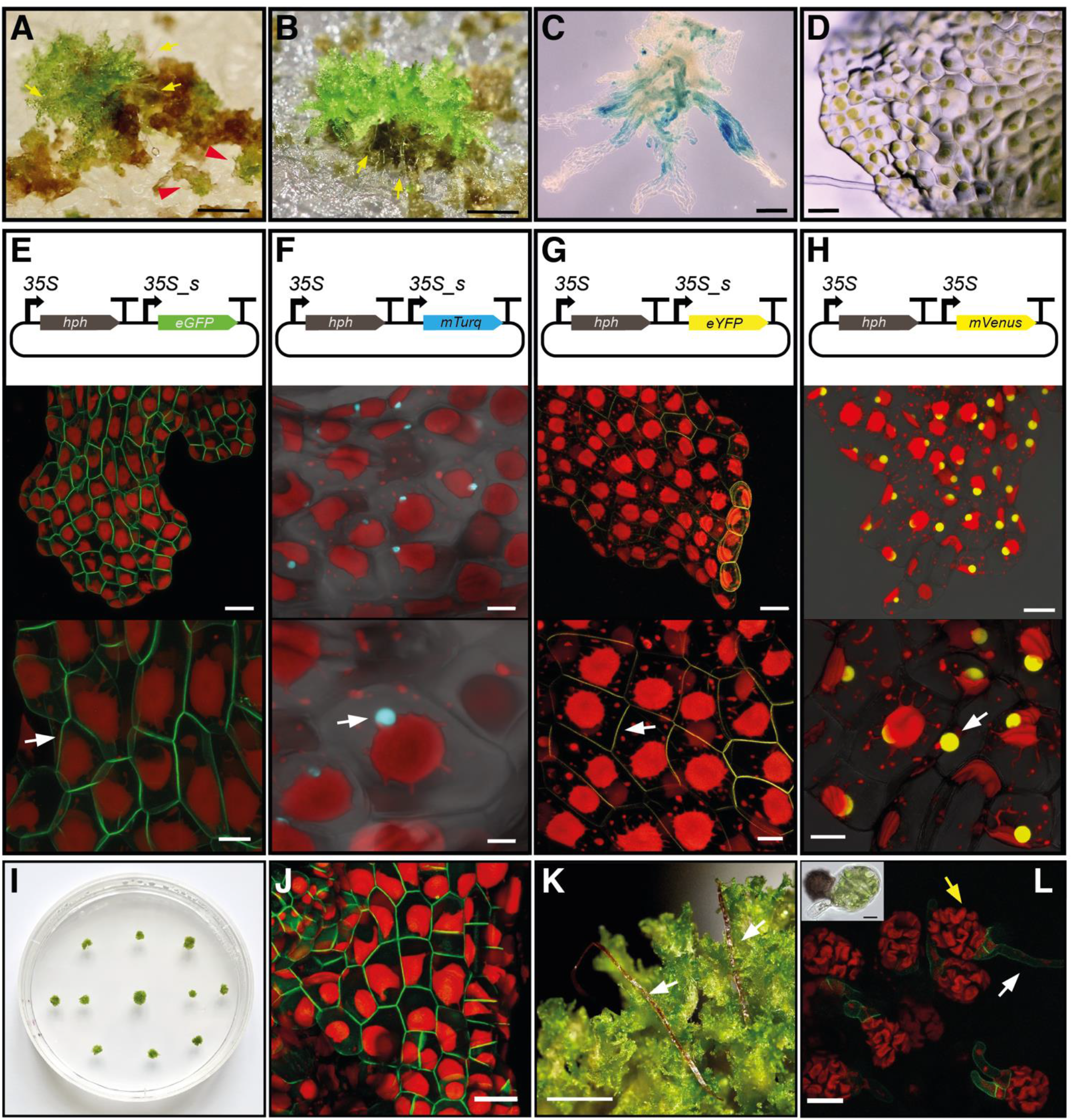
Schematic representation of transformation constructs and transgenic *Anthoceros agrestis* expressing different fluorescent proteins. A-B) The emergence of rhizoids (shown with yellow arrows) is a reliable indicator of successfully transformed plant fragments. In (A) the red arrowheads show false positives regenerating (green) tissue fragments that lack rhizoids. Scale bars: 2 mm. C) GUS activity detected as blue staining in thallus tissue fragments from a plant transformed with the pCAMBIA1305.2 plasmid. Scale bar: 200 μm. D) Light micrograph of thallus surface view of thallus (gametophyte), similar to the area imaged in E-H. Scale bar: 100 μm. E-H top: Schematic representation of constructs for the expression of two transcription units (TU): one TU for the expression of the *hygromycin B phosphotransferase* (*hph*) gene under the control of the cauliflower mosaic virus (CaMV) 35S promoter and one TU for the expression of *p-35S_s::eGFP-LTI6b* (E), *p-35S_s::mTurquoise2-N7* (F), *p-35S_s::eYFP-myr* (G) and *p-35S:mVenus-N7* (H) TU. *hph: hygromycin B phosphotransferase;* 35S: CaMV 35S promoter; eGFP: enhanced green fluorescent protein; mTurquoise2: monomeric turquoise 2 fluorescent protein; eYFP: enhanced yellow fluorescent protein; LTI6b: Low Temperature Induced Protein 6B signal for membrane localization; N7: Arabidopsis At4g19150/N7 nuclear localization signal; myr: myristoylation signal for membrane localization; nosT: 3’ signal of *nopaline synthase*. E-H middle and bottom: Images of *A. agrestis* Oxford thallus tissue expressing different combinations of CaMV 35S promoter - fluorescent protein - localization signal. E) *p-35S_s::eGFP-LTI6b* for plasma membrane localization (white arrow). Scale bars: top: 50 μm, bottom: 20 μm, F) *p-35S_s::mTurquoise2-N7* for nuclear localization (white arrow). Scale bars: top: 20 μm, bottom: 10 μm, G) *p-35S_s::eYFP-myr* for plasma membrane localization (white arrow). Scale bars: top: 50 μm, bottom: 20 μm, H) and *p-35S::mVenus-N7* for nuclear localization (white arrow). Scale bars: top: 50 μm, bottom: 25 μm. The bottom image is a magnification of the image in the middle. Red, chlorophyll autofluorescence. I) Example of transgenic *A. agrestis* plants (gametophyte thallus). Petri dish dimensions: 92 x16 mm. J) Images of *A. agrestis* Bonn gametophyte tissue expressing the *p-35S_s::eGFP-LTI6b* TU for eGFP plasma membrane localization. Scale bar: 50 μm. K) *A. agrestis* Bonn with mature sporophytes indicated with white arrows. Scale bars: 2 mm L) *A. agrestis* Bonn transgenic spores expressing the *p-35S_s::eGFP-LTI6b* TU (white arrow indicates rhizoid and yellow arrow indicates young thallus). Top left: Light microscopy of *A. agrestis* Bonn wild type germinating spore. Scale bar: 20 μm.

### Tests with eGFP as a reporter

Subsequent experiments were carried out using the *A. agrestis* Oxford isolate and a construct containing the enhanced Green Fluorescent Protein (eGFP) reporter gene (Cormack et al., 1996). eGFP makes the identification of successful transformation events easier without the need of laborious GUS staining. For the construction of the eGFP transformation vector, we used the OpenPlant toolkit (Sauret-Gueto et al., 2020) which is based on the Loop assembly Type IIS cloning system (Pollak et al., 2019). All the DNA parts described here are generated following the common syntax (Patron et al., 2015) and are compatible with Type IIS cloning systems, such as GoldenGate and Loop assembly, facilitating the exchange of DNA parts between different laboratories. The transformation vector contained the *hph* gene driven by the 35S promoter and terminated with a 35S polyadenylation signal. It also contained a *p-35S_s::eGFP-Lti6B* transcription unit (same 35S promoter with the one driving *GUSPlus* in pCAMBIA1305.2 plasmid) terminated by the double *nopaline synthase* (*Nos*) - 35S polyadenylation signal (Sauret-Güeto et al., 2020) (Fig. 3E) which was fused to the Low Temperature Induced Protein 6B (Lti6B) signal for membrane localization from *Arabidopsis thaliana* (Arabidopsis) (Cutler et al., 2000). The vector was transformed into *A. agrestis* using the method described above and eGFP was successfully expressed in *A. agrestis* with the expected localization in the plasma membrane (Fig. 3D, E and I). During the course of this study, we generated 157 stable *A. agrestis* transgenic lines expressing the *p-35S_s::eGFP-Lti6B* (Supplemental Table I). There is variability in eGFP expression patterns between different transgenic lines (Supplemental Fig. S8) presumably due to differences in the transgene copy number or the genome location of transgene insertion. A small fraction of hygromycin resistant lines (four out of 157) do not show eGFP fluorescence which could be attributed to potential silencing events or truncation of the inserted T-DNA. These plants have been through at least five rounds of hygromycin antibiotic selection so it is unlikely they are false positives. A total of 15 lines have been propagated vegetatively for more than two years without abolishing transgene expression.

### Testing additional fluorescent proteins, the Bonn isolate and transgene inheritance

Fluorescent proteins have been proven to be a powerful tool for plant cell biology studies, permitting temporal and spatial monitoring of gene expression patterns at a cellular and subcellular level (Berg et al., 2008). In order to expand the palette of fluorescent proteins that can be used in *A. agrestis*, we tested the expression of the monomeric Turquoise 2 fluorescent protein (mTurquoise2) (Kremers et al., 2006; Goedhart et al., 2012), the enhanced yellow fluorescent (eYFP) protein (Orm et al., 1996), and the mVenus fluorescent protein (Kremers et al., 2006). We used a construct similar to the one for the expression of eGFP protein, but with different subcellular localization signals. mTurquoise2 and mVenus were fused to the nuclear-localization peptide sequence of At4g19150/N7 (Cutler et al., 2000) with a linker to the amino (C)-terminus (Cutler et al., 2000), and eYFP was fused to the membrane-targeting myristoylation (myr) signal to the amino (N)-terminus (Resh, 1999). mTurquoise2 (Fig. 2F), eYFP (Fig. 2G) and mVenus (Fig. 2H) were successfully expressed in *A. agrestis* and were targeted to the predicted cellular compartments (for further information on the number of lines generated see Supplemental Table I).

We then tested whether the protocol developed for the Oxford isolate can be used successfully for the Bonn isolate. Four trials resulted in two successful transformants, which is considerably less than the average number of transformants obtained for the Oxford isolate (Fig. 3J).

We finally tested whether the transgene can be stably inherited through the sexual life cycle. Two eGFP expressing transgenic lines for both Oxford and Bonn isolates were brought to sexual reproduction. Sporophytes were produced and young gametophytes germinating from the spores (sporelings) were expressing eGFP indicating that the transgene and its expression was successfully passed on to the next generation (Fig.3K-L).

### Identification and selection of *A. agrestis* endogenous gene promoters

It is important to identify promoters that can be used to drive constitutive transgene expression (i.e. high-level expression across almost all tissues and development stages). Commonly used constitutive promoters in other bryophyte model species include the *M. polymorpha ELONGATION FACTOR 1 ALPHA* (*EF1a*) promoter (Althoff et al., 2014), the rice *Actin1*, and the *M. polymorpha ubiquitin-conjugating enzyme E2* promoter (Sauret-Güeto et al., 2020). Using the genomic sequence and RNA sequencing data (Li et al., 2020), (Fig. 4 A-B) we identified a series of candidate promoter regions as constitutive *A. agrestis* promoters. In particular we selected the promoter regions of the putative *A. agrestis* homologs of *EF1a, Ubiquitin, Actin*, and the Arabidopsis *GAMMA TONOPLAST INTRINSIC PROTEIN* (*Tip1;1*) genes (Fig. 4 A-B, Supplemental Table II). We amplified a 1532 bp long stretch of the 5’ flanking region including the 5’UTR of the *EF1a*, a 933 bp segment for the *Ubiquitin* (*Ubi*), two fragments (1729 bp and 1516 bp) for the *Actin* (that correspond to two different predicted translational start sites), and a 1368 bp putative promoter for the *Tip1;1* gene (Gene models, position of promoters and the corresponding RNA-seq coverage tracks are shown in Supplemental Fig. S9).

**Figure 4.**
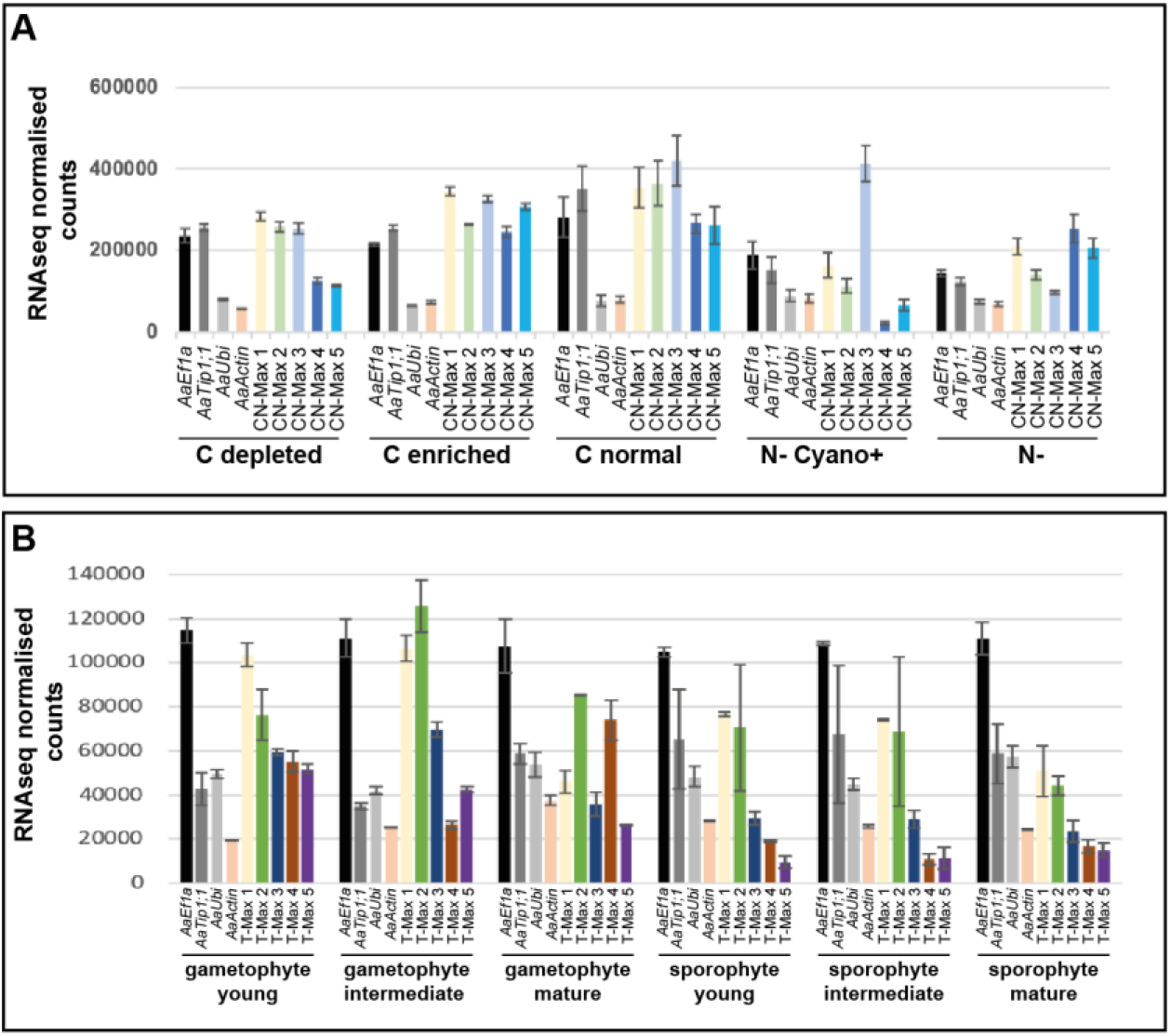
Identification of constitutive promoters for *Anthoceros agrestis*. A) Analysis of expression levels from RNA-seq experiments on *A. agrestis* Oxford using datasets from Li et al. 2020. To generate this dataset, gametophytes were grown under varying carbon sources in the growth medium (indicated as C depleted, C enriched and C normal), as well as two different nitrogen depleted conditions (N-with cyanobacteria symbiosis and N-without cyanobacteria symbiosis). Error bars indicate standard error based on three independent experimental replicates. (Included for comparison: “CN-Max 1, 3 and 5”: highest expressing genes based on N conditions and “CN-Max 2 and 4”: highest expressing genes based on C normal conditions (for gene ID, see Supplemental Table II)). B) Analysis of expression levels from RNA-seq experiments on *A. agrestis* Bonn using data sets from Li et al. 2020. Note: Normalized expression level of *A. agrestis* BONN genes selected to represent strong and constitutive expression across various developmental stages of the gametophyte and the sporophyte phases (Included for comparison: “T-Max 1 to 5”: Highest expressing genes under all conditions (for gene ID, see Supplemental Table II)). Error bars indicate standard error based on two independent experimental replicates.

The candidate promoter regions were cloned (and if necessary domesticated in order to generate a Loop assembly cloning system compatible DNA part), fused with the eGFP or the mTurquoise2 reporter genes, and terminated with the double Nos - 35S terminator (Sauret-Güeto et al., 2020). The *AaEF1a* (Fig. 5A) promoter region was sufficient to drive expression of eGFP throughout the thallus. Similarly, the *AaTip1;1* promoter region was sufficient to drive expression of eGFP and mTurquoise2 (Fig. 5B). However, only three independent lines were obtained for the *p-AaTip1;1::mTurquoise2-N7* construct (with one showing growth retardation probably due to the insertion site of the T-DNA) and one for the *p-AaTip1;1::eGFP* construct (Supplemental Fig. S10). Thus, further characterization of the *AaTip1;1* promoter is needed. The *AaUbi* (Fig. 5C) promoter gave less uniform expression patterns throughout the thallus, and the two *AaActin* promoters produced no detectable signal (Fig. 5D) (summary of the number of lines generated is shown in Supplemental Table I). Our data thus indicate that out of the five candidates, *AaEF1a* is the best promoter for driving relatively strong expression across cells of the gametophyte thallus. We generated a total of eight *p-AaEF1a::eGFP* lines, four of which are shown in Supplemental Fig. S10. Out of the eight hygromycin resistant lines, two do not express eGFP which could be due to transgene silencing or truncation of the inserted T-DNA. In addition, we showed that the *AaEf1a* promoter can drive adequate *hph* expression (Fig. 5E). Finally, we were also able to successfully express simultaneously two different transcription units, *p-AaEF1a::mTurquoise2-N7* and *p-35S_s::eGFP-LTI6b* (Fig. 5F), which was the largest construct (approximately 7.4 kb) we successfully introduced into the *A. agrestis* genome.

**Figure 5.**
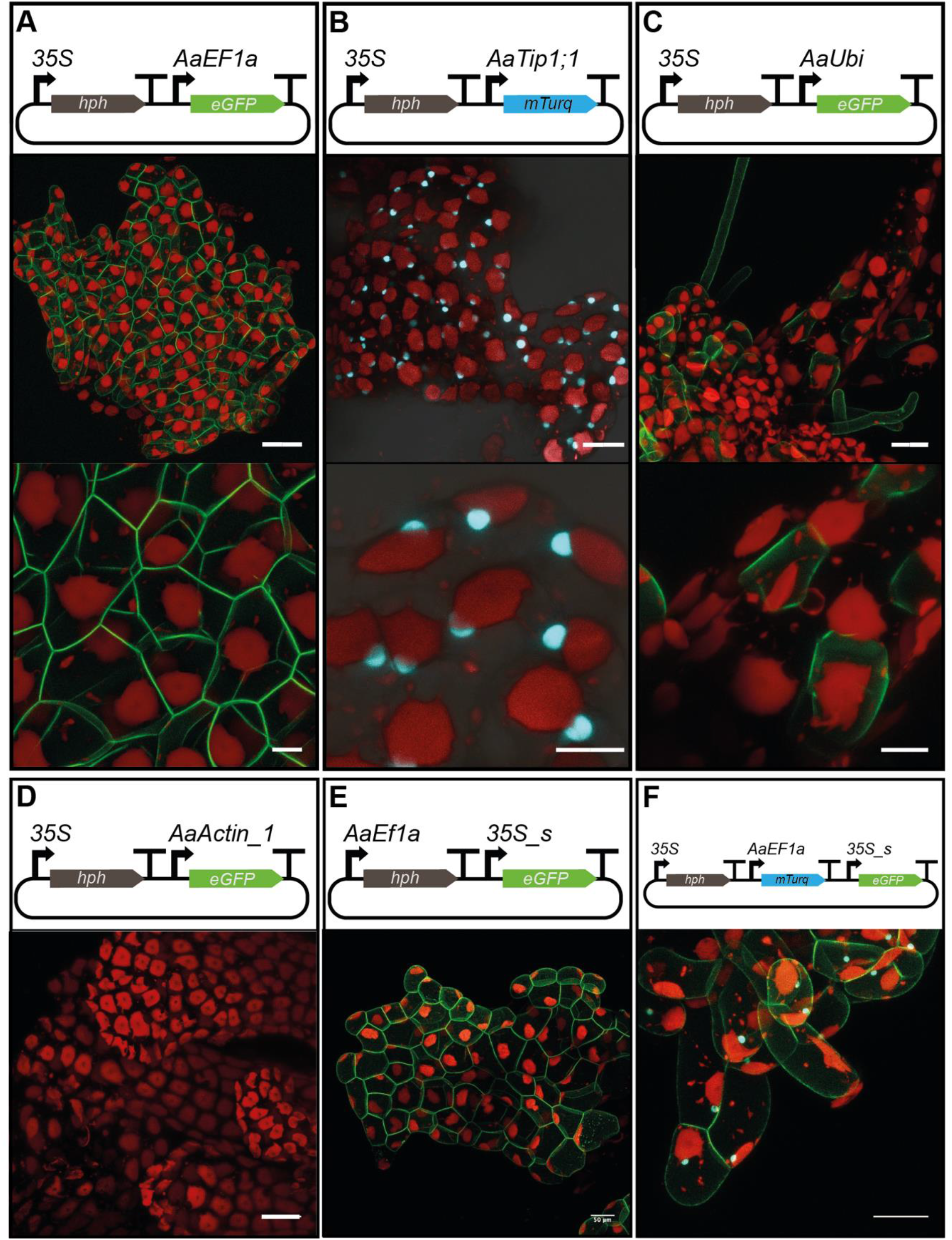
Identification of constitutive promoters for *Anthoceros agrestis*. A-F top: Schematic representation of constructs for the expression of two/three transcription units (TU): one TU for the expression of the *hph* gene under the control of the CaMV 35S or the *p-AaEF1a* promoter and one TU for the expression of plasma membrane localized eGFP and/or nucleus localized mTurquoise2 under the control of different native *A. agrestis* promoters. A) *p-AaEF1a*, B) *p-AaTip1;1*, C) *p-AaUbi* and D) *p-AaActin_1*. E) *p-AaEF1a* driving *hph* and F) *p-AaEF1a* driving *mTurquoise2-N7*. A-F middle and bottom: Images of *A. agrestis* Oxford gametophyte tissue expressing different combinations of *A. agrestis* native promoter - fluorescent protein - localization signal. A) *p-AaEF1a:eGFP-LTI6b* for plasma membrane localization. Scale bars: top: 50 μm, bottom: 10 μm, B) *p-AaTip1;1::mTurquoise2-N7* for nuclear localization. Scale bars: top: 20 μm, bottom: 10 μm, C), *p-AaUbi::eGFP-LTI6b* for plasma membrane localization. Scale bars: top: 20 μm, bottom: 10 μm. The bottom images are a magnification of the images in the top. D) *p-AaActin::eGFP-LTI6b*. Scale bars: 50 μm and E) *p-AaEF1a::hph - p-35S_s::eGFP-LTI6b*. Scale bar: 50 μm. F) Image of *A. agrestis* Oxford gametophyte tissue expressing both *p-AaEF1a::mTurquoise2-N7* for nuclear localization and *p-35S_s::eGFP-LTI6b* for plasma membrane localization. Scale bar: 50 μm. Red, chlorophyll autofluorescence.

### Comparison of the 35S and *AaEF1a* promoters

Expression of eGFP driven by the CaMV 35S promoter seems to be weaker in newly grown parts of the thallus (Fig. 6A and Supplemental Fig. S8). This is similar to the expression patterns of transgenes driven by the CaMV 35S promoter in *M. polymorpha*, which has a strong activity in all parts of the thallus except the notch area (Althoff et al., 2014). Expression of eGFP driven by the *AaEF1a* promoter seems to be stronger in the putatively younger parts of the thallus (Fig. 6B and Supplemental Fig. S10). This is similar to the expression patterns of transgenes driven by the *EF1a* promoter in *M. polymorpha*, showing a strong activity in all parts of the thallus particularly the notch area (Althoff et al., 2014).

**Figure 6:**
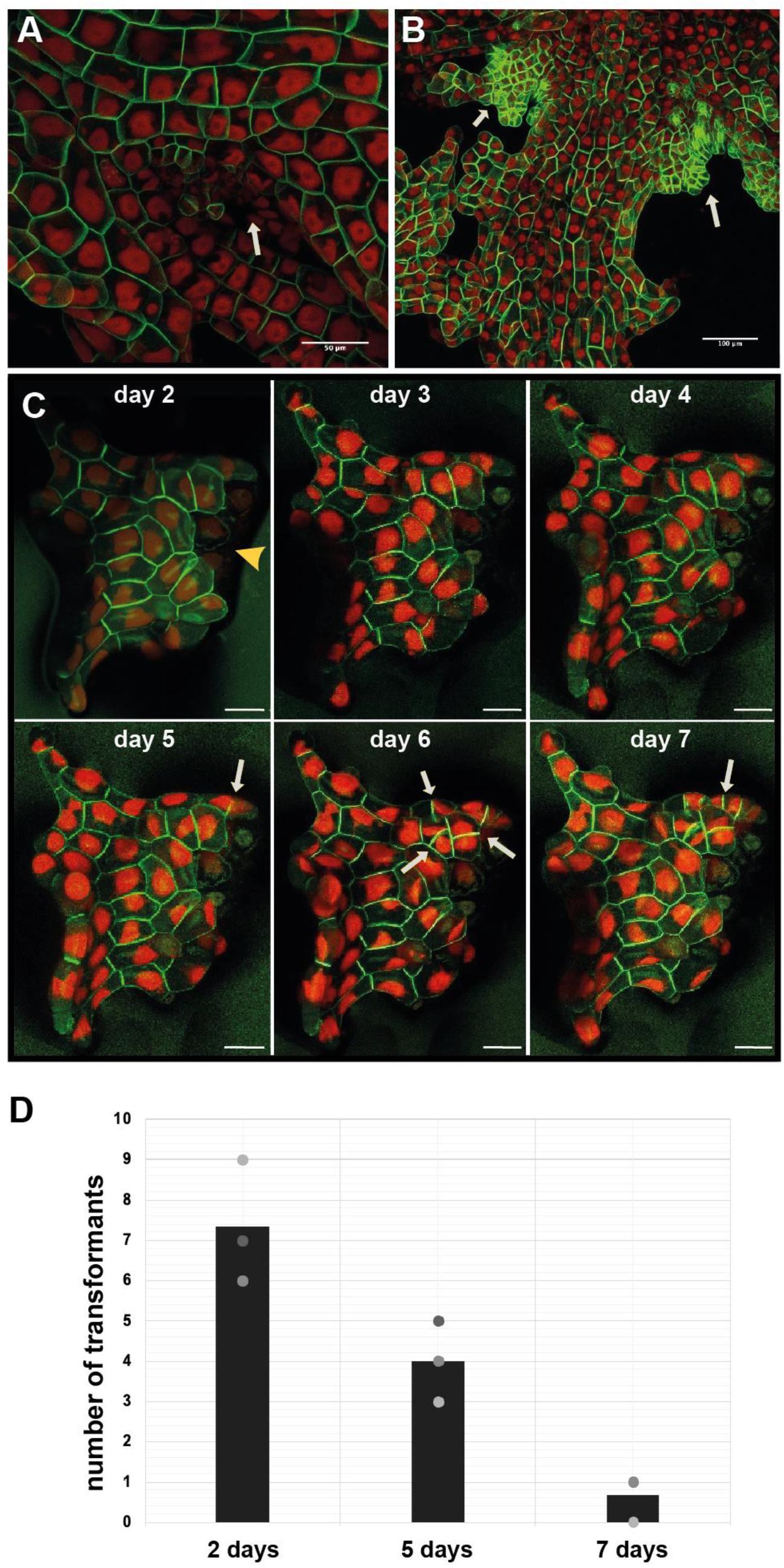
Comparison of the 35S with the *AaEF1a* promoter and factors affecting the efficiency of *Agrobacterium-mediated* transformation of *Anthoceros agrestis*. A) Expression of eGFP driven by the CaMV 35S promoter. Younger part of the thallus indicated with a white arrow. Scale bar: 50 μm. B) Expression of eGFP driven by the *AaEF1a* promoter. Younger part of the thallus indicated with white arrows. Scale bar: 100 μm. C) Confocal microscopy images of fragmented *A. agrestis* thallus tissue taken on seven consecutive days after homogenization. Five days after homogenization plants start to regenerate. Yellow arrowhead indicates the fragmented thallus part. White arrows indicate new cell divisions. Scale bars: 50 μm. D) Number of transgenic lines obtained using *A. agrestis* thallus tissue two, five and seven days after homogenization. Values of independent experimental replicates are shown.

### Transformation efficiency and optimization

In order to estimate the transformation efficiency of the protocol, we performed 10 independent trials using approximately 2 g of tissue as starting material and the *p-35S::hph - p-35S_s::eGFP-LTI6b* construct. The number of successful transformation events per experiments varied from 3 to 23 (Table I).

**Table I:** Number of *p-35S::eGFP-Lti6B* transgenic lines obtained from different transformation experiments.

| **Experiment** | **Amount of tissue (g)** | **Number of transformants** |
| --- | --- | --- |
| **1** | 2.7 | 21 |
| **2** | 3.1 | 12 |
| **3** | 2.9 | 23 |
| **4** | 2.5 | 5 |
| **5** | 1.7 | 3 |
| **6** | 2.7 | 11 |
| **7** | 2.6 | 19 |
| **8** | 1.9 | 4 |
| **9** | 2.23 | 12 |
| **10** | 2.1 | 6 |

We then carried out further experiments to optimize transformation efficiency. We reasoned that tissue susceptibility to *Agrobacterium* infection may differ during different stages of regeneration after homogenization, thereby affecting transformation efficiency. To estimate when plant regeneration is initiated, we set up a microscopy time course using homogenized thallus fragments. The first cell division was observed five days after homogenization (Fig. 6C). Based on this result, we carried out an optimization experiment starting co-cultivation at two, five and seven days after homogenization. We found that the number of stably transformed lines decreased when using tissue that was recovered for more than five days after homogenization (Fig. 6D). The highest number of transformants could be obtained when using tissue two days after homogenization. In parallel with the above mentioned experiment, we also tested another *Agrobacterium* strain, the *GV3101*, for its ability to infect *A. agrestis* thallus. However, only a single successful transformation event was obtained when the *GV3101* strain harboring the *p-35S::hph - p-35S::eGFP-LTI6b* construct was used.

### Verification of transgene incorporation into the *A. agrestis* genome

To confirm the genomic integration of the transgene, we sequenced and assembled the genomes of five stable transformant lines (Fig. 7). For all five lines, we found a single integration site, with one line showing a single full length insertion of the T-DNA. The other four lines additionally showed one or multiple partial insertions in inverted and/or tandem directions (Supplemental Material). To confirm the transgene integration site in the five lines, fragments overlapping the 5’- and 3’-ends of the inserts and their adjacent genomic region were amplified by nested PCR and Sanger-sequenced. The resulting sequences confirmed the integration sites identified by the genome assemblies (Fig. 7G and Supplemental Material). We conclude that the transformation method described here results in the stable integration of one or more targeted transcriptional units into the *A. agrestis* nuclear genome.

**Figure 7:**
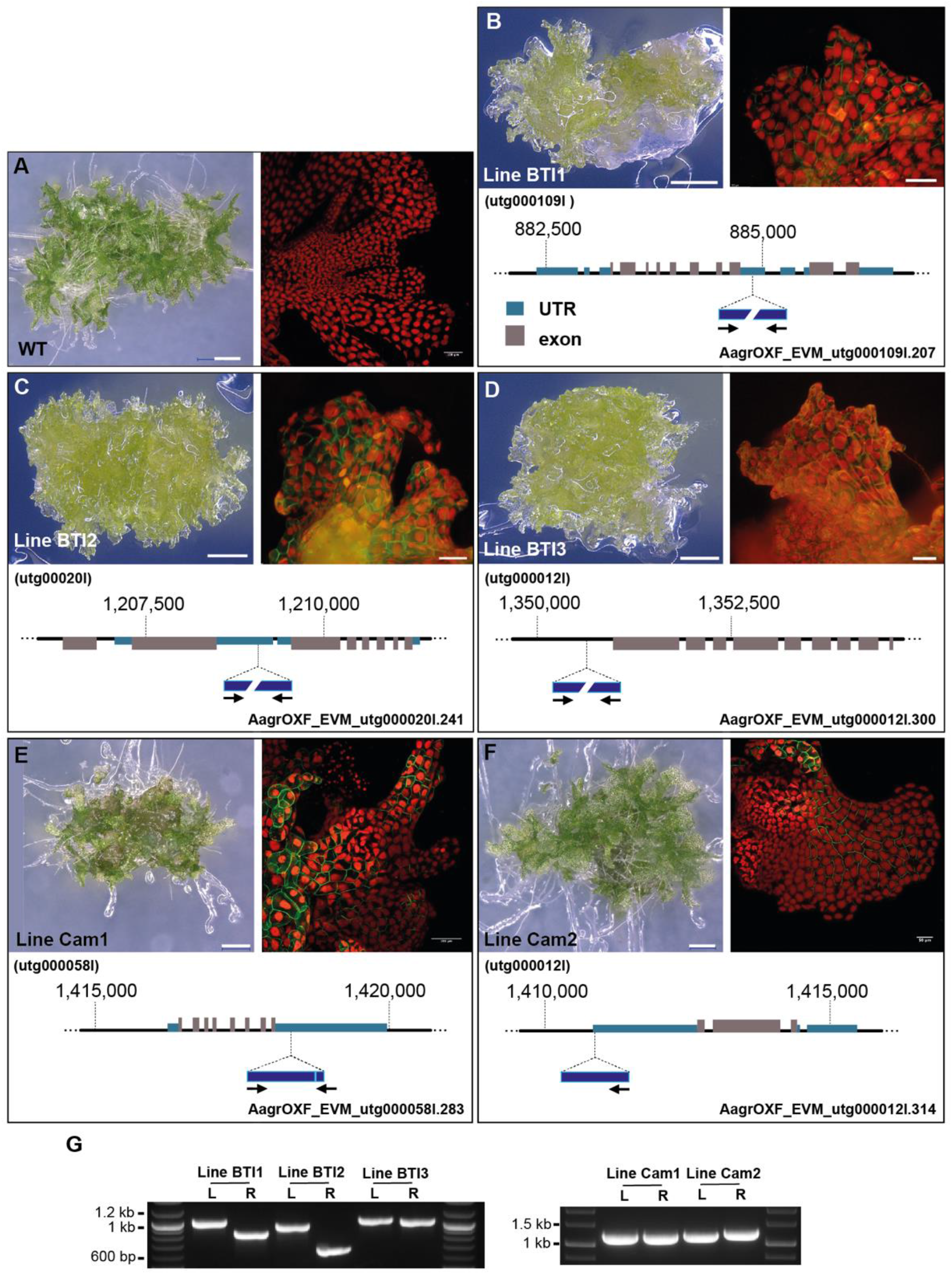
Stable incorporation of transgene into *Anthoceros agrestis* genome. A-F Left: Light micrograph (LM) of transgenic thallus of *A. agrestis* plants. Scale bar: 500 μm. Right: Confocal fluorescent microscopy images of thallus expressing eGFP in the plasma membrane, driven by the CaMV 35S promoter. Scale bars: A-E 100 μm and F 50 μm. Bottom: Location of transgene insertion in the genome (see details in Supplemental material). Black arrows indicate directionality of T-DNA insert. G) PCR analysis of genomic DNA from transgenic plants. L: fragment amplified from sequences spanning the 5’-end of the T-DNA inserts and their respective adjacent genomic regions. R: fragment amplified from sequences spanning the 3’-end of the T-DNA inserts and their respective adjacent genomic regions. Note: A, E and F LM images were acquired using a KEYENCE VHX-S550E microscope (VHX-J20T lens) and confocal fluorescent images with a Leica SP8X microscope, in Cambridge University. B-D LM and fluorescent images were acquired using a Leica M205 FA Stereomicroscope with GFP long-pass (LP) filter, in BT Institute.

## Discussion

The protocol described in this study successfully generated stable transformants in *A. agrestis* Oxford and Bonn isolates and may be applicable to other hornwort species. We generated a total of 216 stable lines. We showed that transgenic lines can be propagated for more than two years without abolishing transgene expression. Additionally, we verified that the transgene is integrated into the genome of *A. agrestis* and can be successfully inherited.

The genome sequencing of transgenic lines showed that the integration occurs in a single locus with one or more copies, which is similar to the reports available for other organisms based on DNA gel blot analysis (Feldmann and David Marks, 1987; Ishizaki et al., 2008; Plackett et al., 2014). The utilization of recent high throughput sequencing technologies combined with the genome size of the *A. agrestis* allows precise determination of the transgene insertion site. Thus, it should be simple to perform enhancer-trap or T-DNA based mutagenesis experiments in *A. agrestis*.

The light conditions under which the tissue was grown significantly affected the thallus morphology and were critical for successful transformation. It is likely that high light intensity triggers the accumulation of secondary metabolites (such as mucilage) and/or affects the composition of the cell wall, thereby significantly reducing transformation efficiency. Multiple photoreceptors are present in the *A. agrestis* genome (Li et al., 2014; Li et al., 2015a; Li et al., 2015b). Identifying which receptors determine the response to high light intensity could help to further improve transformation efficiency. In addition, testing different methods for tissue fragmentation, such as vortexing or use of a scalpel, or employing other types of tissues (germinating spores or callus) might also improve transformation efficiency.

We are currently developing genome editing tools for *A. agrestis* using CRISPR/Cas9 (Jinek et al., 2012). We are also investigating whether inducible gene expression systems such as the glucocorticoid receptor (Schena et al., 1991) or estrogen receptor (Zuo et al., 2000) can be applied successfully in hornworts. Finally, we are testing alternative gene delivery methods for both *A. agrestis* Oxford and Bonn isolates, such as particle bombardment.

The development of a hornwort transformation method, in combination with the recently published genome, will greatly facilitate more comprehensive studies of the mechanisms underpinning land plant evolution. It can also help with engineering hornwort traits into plants with agronomic value. For example engineering pyrenoids in crops, has the potential toimprove carbon fixation and therefore increase crop yield (Li et al., 2017).

## Materials and methods

### Plant material and maintenance

The *Anthoceros agrestis* Oxford and Bonn isolates were used (Szövényi et al., 2015). *A. agrestis* thallus tissue was propagated on KNOP medium (0.25 g/L KH_2_PO_4_, 0.25 g/L KCl, 0.25 g/L MgSO_4_•7H_2_O, 1 g/L Ca(NO_3_)2•4H_2_O and 12.5 mg/L FeSO_4_•7H_2_O). The medium was adjusted to pH 5.8 with KOH and solidified using 7.5 g/L Gelzan CM (#G1910, SIGMA) in 92×16 mm petri dishes (#82.1473.001, SARSTEDT) with 25-30 mL per plate. Plants were routinely grown in a tissue culture room (21°C, 12 h of light and 12 h of dark, 5 μmol m^-2^ s^-1^ light intensity). In order to subculture the thallus tissue, a small part of the thallus was cut using sterile disposable scalpels (#0501, Swann Morton) and placed on fresh media on a monthly basis.

### Co-cultivation medium

Co-cultivation medium was liquid KNOP supplemented with 2% sucrose (0.25 g/L KH_2_PO_4_, 0.25 g/L KCl, 0.25 g/L MgSO_4_•7H_2_O, 1 g/L Ca(NO_3_)2•4H_2_O, 12.5 mg/L FeSO_4_•7H_2_O and 20 g/L sucrose, pH 5.8 adjusted with KOH).

### Tissue preparation for transformation

Approximately 2 g of thallus tissue were divided into 4 parts, and each part was homogenized in 15 mL of sterile water using a homogenizer (#727407, IKA Ultra-Turrax T25 S7 Homogenizer) and corresponding dispensing tools (#10442743, IKA Dispersing Element), for 5 seconds, using the lowest speed of 8000 rpm. The homogenized tissue was washed with 50 mL of sterile water using a 100 μm cell strainer (#352360, CORNING), spread on solid KNOP medium and placed at 21°C under 12 hours light and 12 hours dark at a light intensity of 3-5 μmol m^-2^ s^-1^. After 4 weeks the tissue was re-homogenized in 15-20 mL of sterile water and filtered using a 100 μm cell strainer. The re-homogenized tissue was transferred again onto 4 plates with solid KNOP medium and was allowed to grow for 2 days at 21°C under continuous light (35 μmol m^-2^ s^-1^, PHILIPS, TL-D58W/835).

### *Agrobacterium* culture preparation

A single *Agrobacterium* colony (AGL1 strain) was inoculated in 5 mL of LB medium supplemented with rifampicin 10 μg/mL (#R0146, Duchefa), carbenicillin 50 μg/mL (#C0109, MELFORD) and the plasmid-specific selection antibiotic spectinomycin 100 μg/mL (#SB0901, Bio Basic). The pre-culture was incubated at 28°C for 2 days with shaking at 120 rpm. OD600 was ~2.7 and was measured using an OD600 DiluPhotometer (IMPLEN).

### Co-cultivation conditions

5 mL of 2 days *Agrobacterium* culture was centrifuged for 7 minutes at 2000 xg. The supernatant was discarded and the pellet was resuspended in 5 mL liquid KNOP supplemented with 2% (w/v) sucrose (#S/8600/60, Fisher) and 100 μM 3’,5’-dimethoxy-4’-hydroxyacetophenone (acetosyringone) (#115540050, Acros Organics, dissolved in dimethyl sulfoxide (DMSO) (#D8418, SIGMA)). The culture was incubated with shaking (120 rpm) at 28°C for 5 hours. The regenerating thallus tissue was transferred (1/2 tissue from one plate – 2 days after the second homogenization) into a well of a 6-well plate with 4 mL of liquid KNOP medium supplemented with 2% (w/v) sucrose. 80 μL of *Agrobacterium* culture and acetosyringone at final concentration of 100 μM were added to the medium.

The tissue and *Agrobacterium* were co-cultivated using a 6-well plate (#140675, ThermoFisher) for 3 days with shaking at 110 rpm at 22°C on a shaker without any additional light supplemented (only ambient light from the room, 1-3 μmol m^-2^ s^-1^). After 3 days, the tissue was drained using a 100 μm cell strainer (#352360, CORNING) and moved onto solid KNOP medium plates supplemented with 100 μg/mL cefotaxime (#BIC0111, Apollo Scientific) and 10 μg/mL Hygromycin (#10687010, Invitrogen). After 3-4 weeks, plants were transferred to fresh solid KNOP medium plates supplemented with 100 μg/mL cefotaxime and 10 μg/mL Hygromycin and grown at 22°C 12 hours light and 12 hours dark at a light intensity of 35 μmol m^-2^ s^-1^ (PHILIPS, TL-D58W/835).

### GUS staining

GUS staining was performed according to Plackett et al., 2014.

### Genomic DNA extraction

A modified CTAB protocol from (Porebski et al., 1997) was used for hornwort genomic DNA extraction. 0.5 g of tissue was harvested and frozen in liquid nitrogen. Tissue was ground into a fine powder using a chilled mortar and pestle and then added to 10 mL of DNA extraction buffer (100 mM Tris-HCl pH 8, 1.4 M NaCl, 20 mM EDTA pH 8, 2% (w/v) CTAB, 0.3% (v/v) β-mercaptoethanol and 100 mg of polyvinylpyrrolidone (PVP)/g of tissue) that had been prewarmed at 60°C, 100 μL of RNAse A (100 mg/mL) was added and the solution was mixed well. The mix was incubated at 60°C for 20-30 minutes and then removed from heat and allowed to cool at room temperature for 4 minutes. 12 mL of chloroform:isoamyl alcohol (24:1) was added, mixed well by inversion and then centrifuged at 12000 xg for 10 minutes at room temperature. The upper aqueous phase was transferred to a new 50 mL centrifugation tube and 10 mL of chloroform:isoamyl alcohol (24:1) was added, mixed well by inversion and then centrifuged at 6000 xg for 10 minutes at room temperature to remove any remaining PVP in the aqueous phase. The upper aqueous phase was transferred to a new 50 mL centrifugation tube and ½ volume of 5 M NaCl was added. 2 volumes of cold (−20°C) 95% (v/v) ethanol were also added and the contents of the tube were mixed well by inversion. The tube was spun at 20000 xg for 6 minutes. The pellet was resuspended in 2 mL of TE buffer and the previous step was repeated. The pellet was washed with cold 70% (v/v) ethanol. The pellet was dried and dissolved in 80 μL of TE buffer and then stored at 4°C.

### Construct generation

Constructs were generated using the OpenPlant toolkit (Sauret-Gueto et al., 2020). Full sequence of constructs can be found in Supplemental Table III.

### Promoter identification and isolation

We used RNA-seq data to find genes showing constantly high levels of expression under various developmental stages and experimental conditions (“constitutively expressed genes”). To do so, we estimated expression of genes under three developmental stages of the gametophyte and sporophyte phases and in symbiosis with cyanobacteria. We retrieved raw RNA-seq data for these experiments from (Li et al., 2020). We used trimmomatic to quality filter and trim the raw reads. Gene expression was estimated using Salmon (Patro et al., 2017) and expressed as normalized expression counts. We identified candidates by selecting those showing the highest average expression level and the least gene expression variability across all conditions investigated. We then manually selected a subset of genes taking into account their genomic location, exact expression pattern, and the length and sequence composition of their putative promoter sequences. We also assessed the suitability of our candidate promoters using the information available for *M. polymorpha* and *P. patens* (Supplemental Table II).

Putative promoter sequences were amplified from genomic DNA using the KOD Hot start polymerase (#71086-5, Merck Millipore) and cloned into pJET1.2 (#K1231, ThermoFisher) before Sanger sequencing. Loop assembly compatible DNA parts were generated according to (Sauret-Güeto et al., 2020). List of primers can be found in Supplemental Table IV.

### Sample preparation for Imaging

A gene frame (#AB0576, ThermoFisher) was positioned on a glass slide and 30 μL of KNOP medium with 1% (w/v) Gelzan CM (#G1910, SIGMA) was placed within the gene frame. A thallus fragment was placed within the medium-filled gene frame together with 30 μL of milliQ water. The frame was then sealed with a cover slip. Plants were imaged immediately using a Leica SP8X spectral fluorescent confocal microscope.

For the regeneration test experiment, five thallus fragments were placed into a KNOP medium-filled gene frame as described above (three slides and 15 plants in total). Images were acquired on a daily basis, for a total duration of a week, using a Leica SP8X spectral fluorescent confocal microscope and a 10× air objective (HC PL APO 10×/0.40 CS2).

### Imaging with Confocal Microscopy

Images were acquired on a Leica SP8X spectral confocal microscope. Imaging was conducted using either a 10× air objective (HC PL APO 10×/0.40 CS2) or a 20× air objective (HC PL APO 20×/0.75 CS2). Excitation laser wavelength and captured emitted fluorescence wavelength window were as follows: for mTurquoise2 (442 nm, 460–485 nm), for eGFP (488 nm, 498-516 nm), for mVenus and eYFP (515 nm, 522-540 nm), and for chlorophyll autofluorescence (488 or 515, 670-700 nm). When observing lines expressing both eGFP and mTurquoise2, sequential scanning mode was used.

### Light microscopy

Images were captured using a KEYENCE VHX-S550E microscope (VHX-J20T lens) or a Leica M205 FA Stereomicroscope (with GFP longpass (LP) filter).

### Sequencing transformant line genomes

Transformant lines were grown on solid KNOP medium containing 10 μg/mL hygromycin and 100 μg/mL cefotaxime. Genomic DNA was extracted from 600 mg fresh tissue per line using either the DNeasy Plant Pro kit (#69204, Qiagen) (Cam-1 and Cam-2 lines) or the procedure from Li et al (2020) (BTI1-3 lines) to reach a total yield of at least 200 ng/line. Illumina libraries were prepared using the TruSeq DNA nano kit (#20015964, Illumina) and were sequenced on an Illumina Novaseq 6000 platform with an expected sequencing depth of 80-150x for all five lines in paired-end mode (read length: 151 bp).

After sequencing we quality filtered and trimmed reads using trimmomatic (command line: ALL_TruSeq-PE.fa:2:30:10:2:keepBothReads LEADING:3 TRAILING:3 SLIDINGWINDOW:4:20 MINLEN:36) and assembled the reads with spades3.14.1 using the --isolate --cov-cutoff auto – only-assembler options as recommended (Nurk et al., 2013). To localize the insertion and its copy number, we used the insert sequence as a query in a BLASTN search (Altschul et al., 1990) against the database containing the assembly (Altschul et al., 1990) with an e-value threshold of 10^-4^. As evidence of genomic integration, we only accepted hits covering the full-length of the insert sequence with one or no mismatches. We manually inspected blast hits to eliminate false positives. Finally, we used the *A. agrestis* Oxford genome sequence (Li et al., 2020) to localize the hits on the pseudomolecules.

### PCR confirmation of transformant insertion sites

Based on the genome assemblies of the transformant lines, primers were designed to amplify regions of 0.6-1.3 kb spanning the 5’- and 3’-end of the T-DNA inserts and their respective adjacent genomic regions. Sequences were amplified from genomic DNA with Phusion High-Fidelity DNA Polymerase (#F-530S, ThermoFisher). For each amplified region, a second set of nested primers were designed to amplify a shorter amplicon, using 1:100 dilution of the previous PCR product as a template. Resulting nested PCR-products were either purified and Sanger sequenced (Eurofins) from both ends, or cloned into pJET1.2 (#K1231, ThermoFisher) before Sanger sequencing from both ends using pJET1.2 sequencing primers.

## Supporting information

All Supp. Figs and Tabs excluding Supp. Tab. 3

## Author Contributions

KS, EF, HT, MW and PS, conceived and designed the experiments. TN and MW identified gene promoter regions. EF and MW performed cloning. EF generated and characterized the transgenic lines and performed imaging. XX, MT and JVC generated the BTI lines and AG imaged the lines. MW, PS, FW and AG analyzed the transcriptomic data. MW performed NGS of transgenic lines, PS and MW assembled the genomes. AG and MW confirmed the insert locations. YY provided technical assistance. KS, EF, TN, MW and PS wrote the article with contributions from all the authors.

## Acknowledgments

We would like to thank Juan Carlos Villarreal for introducing us to the world of hornworts and always generously sharing his expertise. We are also thankful to Jim Haseloff for support during the latest stage of the project. Figure 1D picture credit: John Baker, Oxford University. This research was supported by the Japanese Society for the Promotion of Science (JSPS) Short Term Postdoctoral Fellowship grant no. PE14780 to E.F., the UZH Forschungskredit Candoc grant no. FK-19-089 to M.W., the MEXT and JSPS Grants-in-Aid for Scientific Research on Innovation Areas nos. 25113001 and 19H05672 to H.S., JSPS KAKENHI 26650143 and 18K06367 to K.S., 15H04413 to T.N., 19K22448 to T.N. and K.S., the Swiss National Science Foundation (grants 160004, 131726 and 184826) to P.S., The Deutsche Forschungsgemeinschaft (DFG - German Research Foundation) under the Priority Programme “MAdLand - Molecular Adaptation to Land: Plant Evolution to Change” (SPP 2237, 440370263) to P.S., The Georges and Antoine Claraz Foundation to P.S., M.W and Y.Y, The University Research Priority Program ‘Evolution in Action’ of the University of Zurich to P.S., and the National Science Foundation IOS-1923011 to F.-W.L. and J.V.E.

## Accession numbers

Raw sequencing reads have been submitted to the NCBI SRA under the BioProject ID PRJNA683066 (SRR13209765-SRR13209769).

