## Supplementary material for "A simple *Agrobacterium*-mediated stable transformation technique for the hornwort model *Anthoceros agrestis*": All Supp. Figs and Tabs excluding Supp. Tab. 3

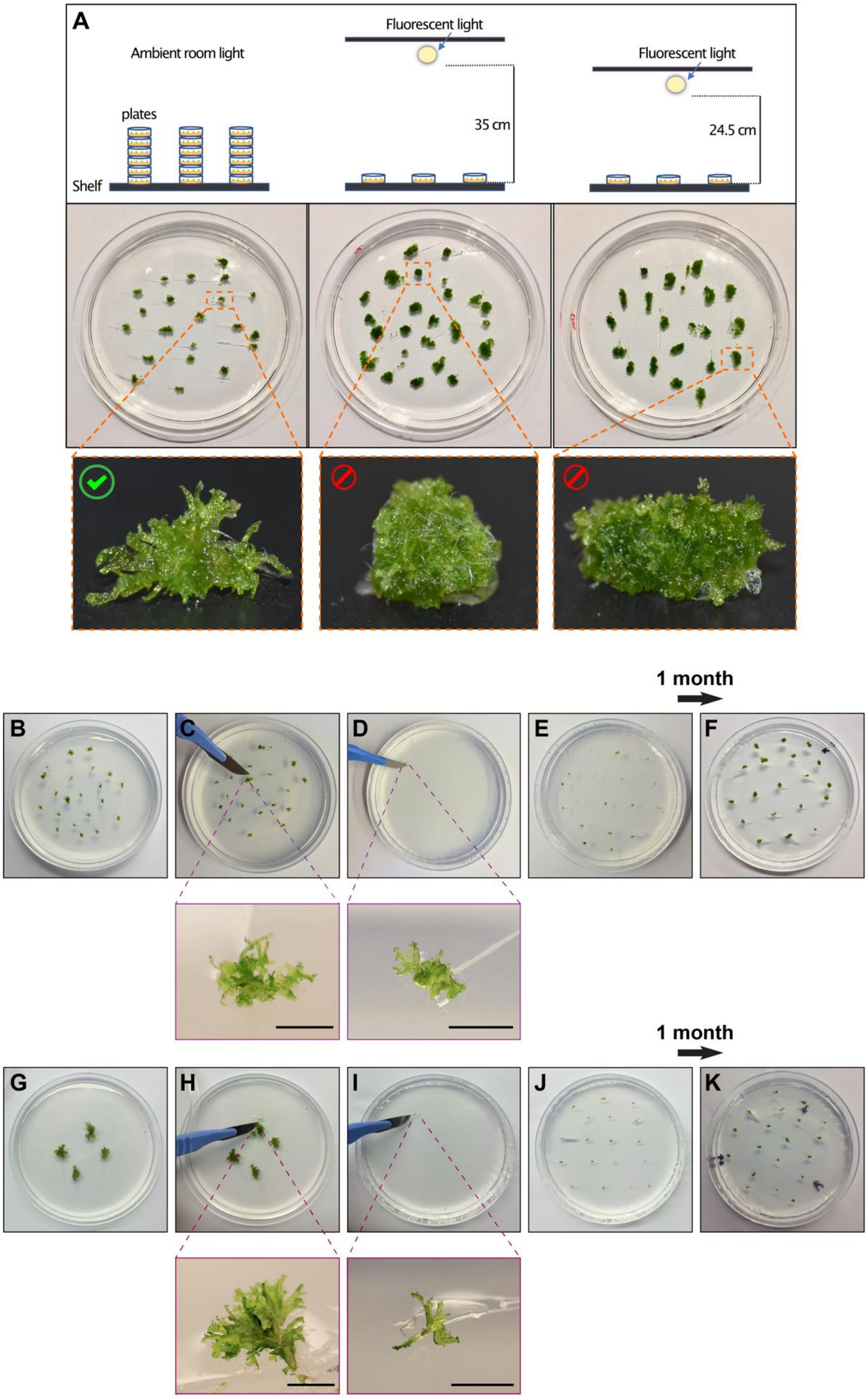
A
Ambient room light
Fluorescent light
35 cm
Fluorescent light
24.5 cm
plates
Shelf
1 month
B
C
D
E
F
1 month
G
H
I
J
K

**Supplemental Figure S1: Effect of light on *A. agrestis* growth and tissue culturing**

A) Examples of *A. agrestis* tissue grown under different light regimes. From left to right: i) tissue morphology when plates were stacked in the laboratory under ambient room light (3-5 μmol $m^{-2}$ $s^{-1}$), ii) tissue morphology under light supplemented by fluorescent tubes, 35 μmol $m^{-2}$ $s^{-1}$, light intensity (PHILIPS, TL-D58W/835) and iii) tissue morphology under light supplemented by fluorescent tubes, 20 μmol $m^{-2}$ $s^{-1}$ light intensity (PHILIPS, TL-D 36W/840). Tissue similar to (i) is optimal for transformation. Use of tissue similar to (ii) and (iii) should be avoided.

B-K) Axenic cultures of *A. agrestis* Oxford (B-F) and Bonn (G-K) gametophytes can be routinely propagated by monthly sub-culturing. For sub-culturing, a small piece of thallus tissue is cut using sterile disposable scalpels and placed on plates containing fresh growth medium. Scale bars: 2 mm. Petri dish dimensions: 92 x16 mm.

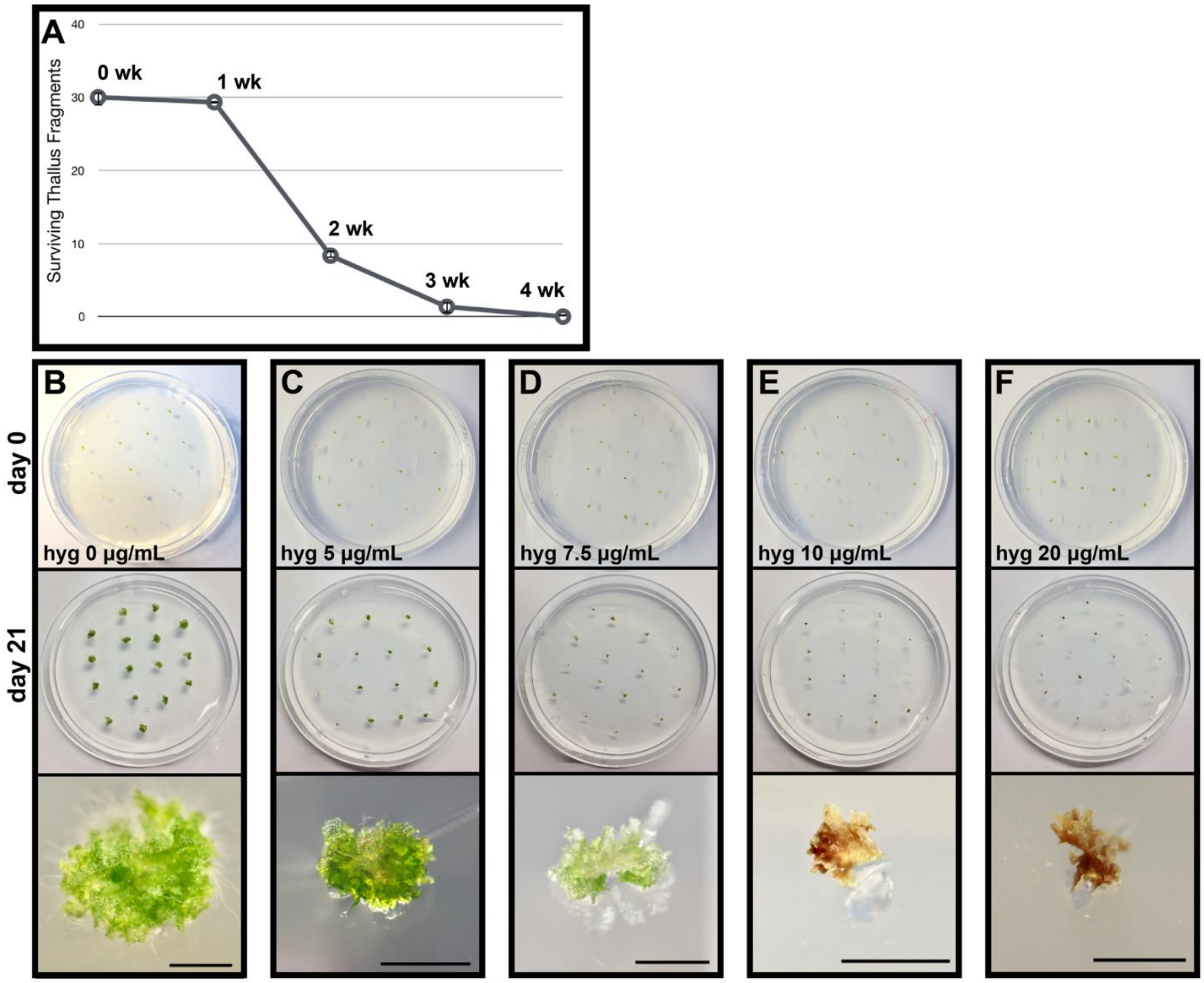


**Supplemental Figure S2. Hygromycin sensitivity of *A. agrestis* gametophytes (Oxford)**

A) Frequency of *A. agrestis* Oxford thallus survival after incubation on KNOP media containing 0 - 20 µg/mL hygromycin B. Values are the mean of three independent replicates for each treatment combination (0 - 20 µg/mL hygromycin selection applied for 0, 1, 2, 3 and 4 weeks). Each replicate comprised 30 fragments of thallus. Error bars indicate standard error based on three independent experimental replicates. Hygromycin treatment killed all thallus fragments within 4 weeks. B-F) Petri dishes with *A. agrestis* Oxford thallus subjected to 0 - 20 µg/mL hygromycin antibiotic selection at day 0 of application of selection (top), day 21 (middle) and magnified images of tissue on day 21 (bottom). Scale bars: 2mm. Petri dish dimensions: 92 x16 mm.

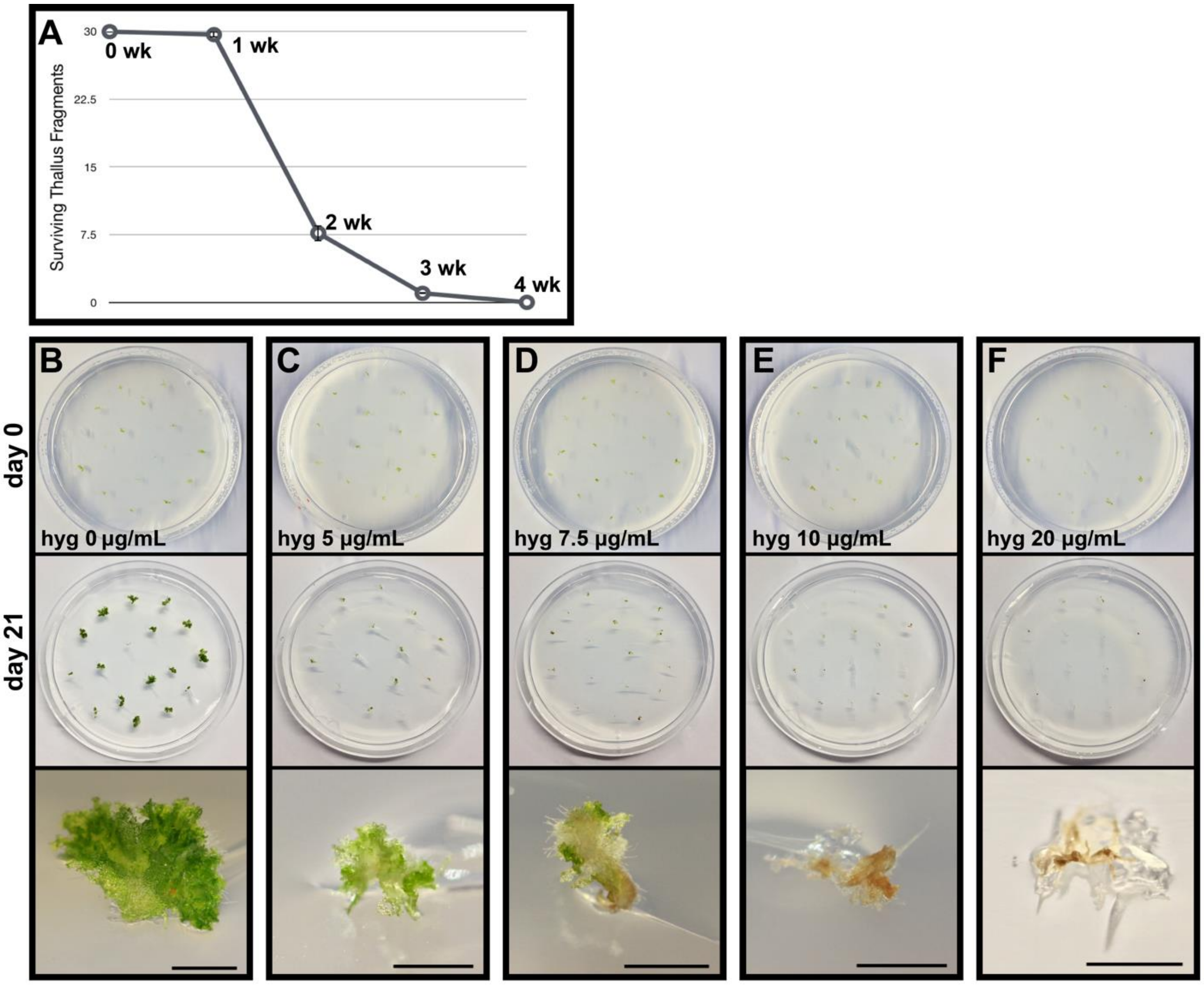


**Supplemental Figure S3. Hygromycin sensitivity of *A. agrestis* gametophytes (Bonn)**

A) Frequency of *A. agrestis* Bonn thallus survival after incubation on KNOP media containing 0 - 20 µg/mL hygromycin B. Values are the mean of three independent replicates for each treatment combination (0 - 20 µg/mL hygromycin selection applied for 0, 1, 2, 3 and 4 weeks). Each replicate comprised 30 fragments of thallus. Error bars indicate standard error based on three independent experimental replicates. Hygromycin treatment killed all thallus fragments within 4 weeks. B-F) Plates with *A. agrestis* Bonn thallus subjected to 0 - 20 µg/mL hygromycin antibiotic selection at day 0 of application of selection (top), day 21 (middle) and magnified images of tissue on day 21 (bottom). Scale bars: 2mm. Petri dish dimensions: 92 x16 mm.

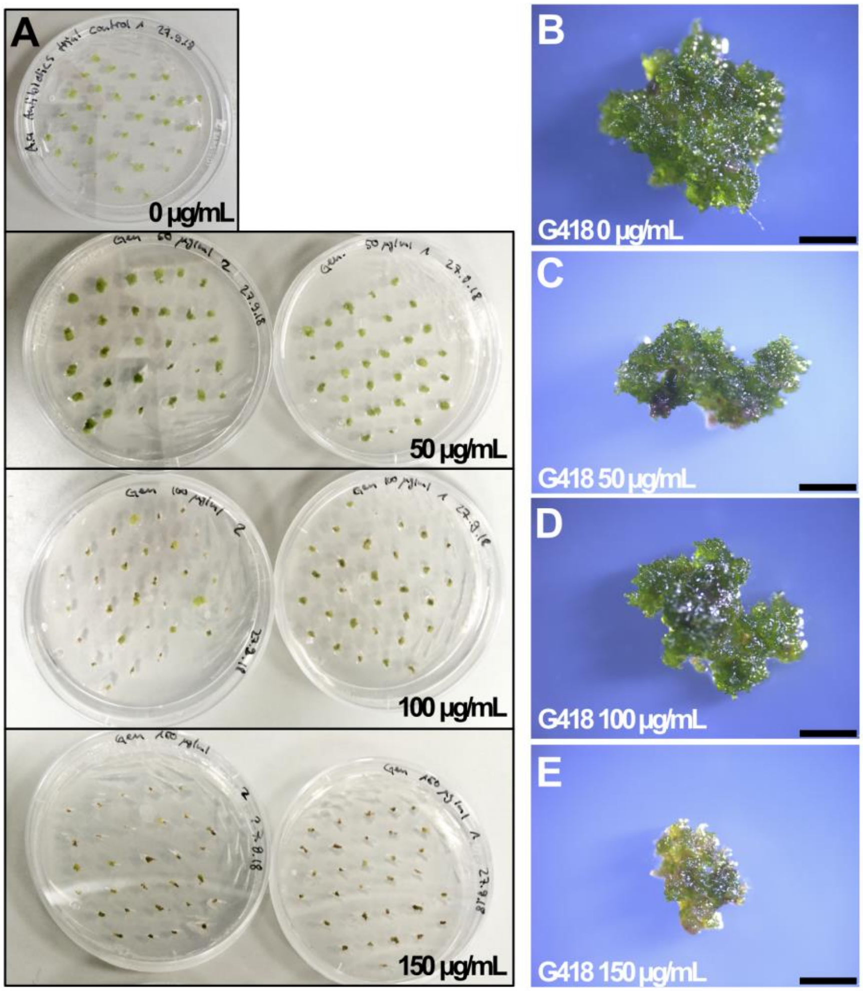


**Supplemental Figure S4. G418 sensitivity of *A. agrestis* gametophytes (Oxford)**

A) Plates with *A. agrestis* Oxford thallus subjected to 0 - 150 µg/mL G418 (#G64000, MELFORD) antibiotic selection for 4 weeks. B-E) magnified images of tissue subjected to 0 - 150 µg/mL G418 antibiotic selection for 4 weeks . Petri dish dimensions: 92 x16 mm.

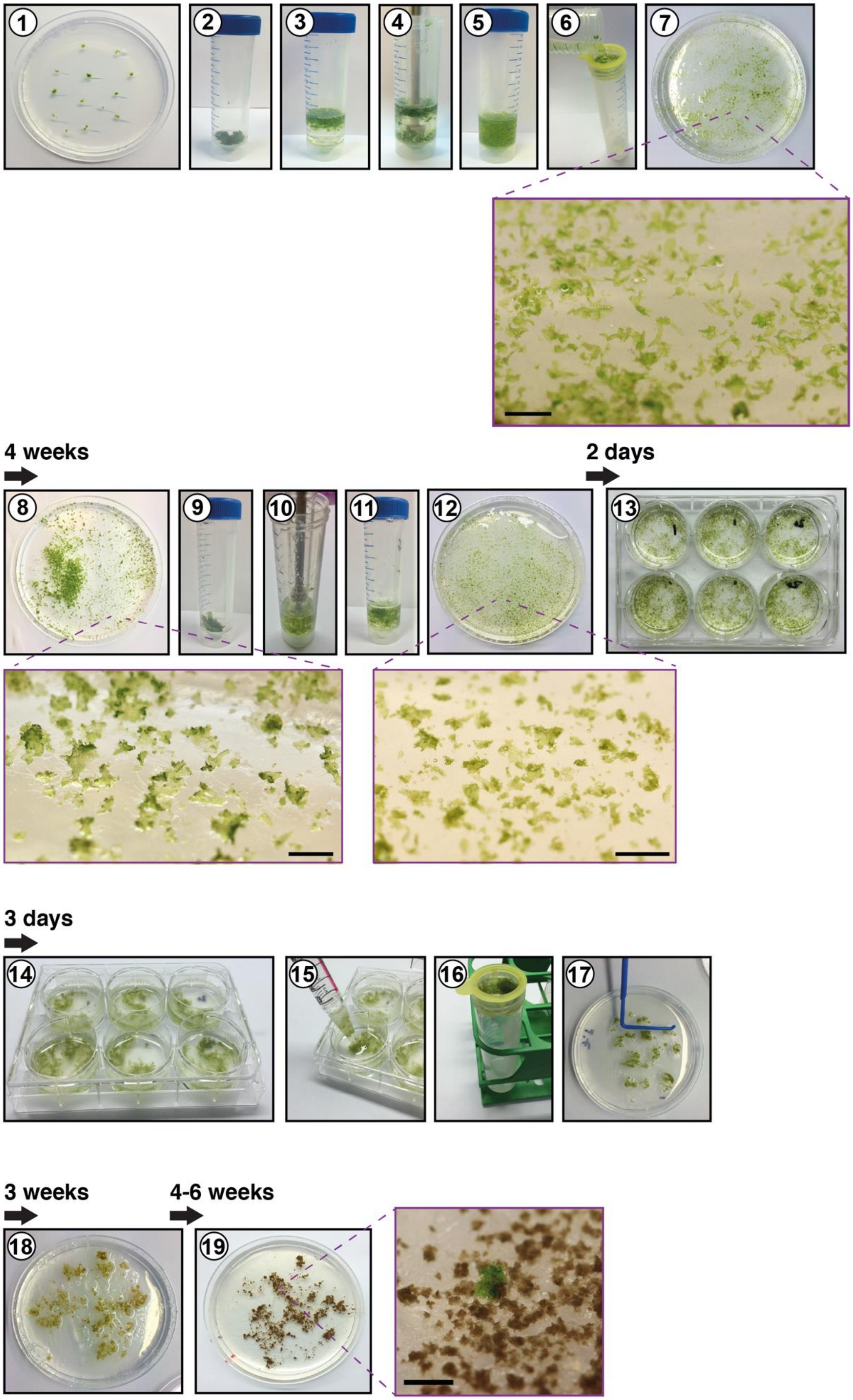
1
2
3
4
5
6
7
4 weeks
8
9
10
11
12
2 days
13
3 days
14
15
16
17
3 weeks
18
4-6 weeks
19

**Supplemental Figure S5: *A. agrestis* Oxford transformation protocol**

**1)** Approximately 2 g of thallus tissue grown for 4 weeks under low light intensity were collected (approximately 0.1 g of tissue per petri dish - 20 petri dishes in total). **2-5)** Tissue was split into three parts, transferred into three 50 mL falcon tubes containing 15 mL of sterile water and homogenized using a homogenizer and corresponding dispensing tools (for 5 sec, lowest power 8000 rpm). **6-7)** The homogenized tissue from one falcon tube was transferred into a cell strainer, was washed with 50 mL of sterile water and transferred onto two petri dishes (six plates in total) containing solid KNOP medium and placed at 21°C under 12 hours light and 12 hours dark, and light intensity 3-5 μmol $m^{-2}$ $s^{-1}$. **8-9)** After 3 weeks the tissue was transferred from the petri dishes into a falcon tube using sterile scalpels, **10-11)** re-homogenized in 15-20 mL of sterile water and filtered using a cell strainer **12)** spread out again onto 4 petri dishes with solid KNOP medium (if necessary, to facilitate spreading of the tissue, 2 mL of sterile water was added to the petri dish) and was allowed to grow for 2 days at 21°C under continuous light (intensity 30 μmol $m^{-2}$ $s^{-1}$). 5 mL of 2 days *Agrobacterium* culture was centrifuged for 7 min at 1800 xg. **13)** The regenerating thallus tissue was transferred into a well of a 6-well plate (transfer ½ the tissue from one plate into a single well) with 4 mL of liquid KNOP medium supplemented with 2% (w/v) sucrose. 80 μL of *Agrobacterium* culture and acetosyringone at a final concentration of 100 μM were added to the liquid medium. **14)** The tissue was co-cultivated with the *Agrobacterium* for 3 days on a shaker at 110 rpm, with only ambient light (1-3 μmol $m^{-2}$ $s^{-1}$). **15-17)** Using a sterile plastic pipette the tissue from a single well was transferred into a cell strainer, drained and then transferred on growth media containing the appropriate antibiotic. To facilitate spreading of the tissue, 2 mL of sterile water was added to the petri dish. **18)** After 3 weeks the tissue was transferred to fresh growth media containing appropriate antibiotics. To facilitate spreading of tissue on the petri dish 2 mL of sterile water was added. **19)** After 6-8 weeks successful transformants were visible on the petri dish (successful transformants can be identified using a microscope after 4 weeks selection based on rhizoid production and/or fluorescence). To eliminate false positives, surviving tissue fragments were transferred again on antibiotic containing growth media.

Petri dish dimensions: 92 x16 mm.

Scale bars: 2 mm.

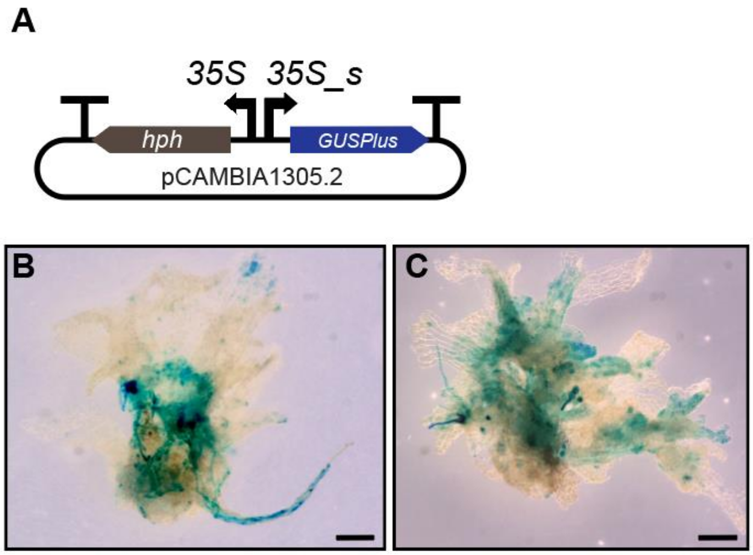


**Supplementary Figure S6: GUS activity detected in *A. agrestis* plants transformed with the pCAMBIA1305.2 plasmid**

A) Schematic representation of the pCAMBIA1305.2 plasmid which contains a cassette for the expression of an intron (catalase) containing GUS gene (GUSPlus) under the control of the CaMV 35S promoter. B-C) GUS activity detected as blue staining in thallus tissue fragments from 2 independent lines (B and C) transformed with the pCAMBIA1305.2 plasmid. Scale bars: 200 μm.

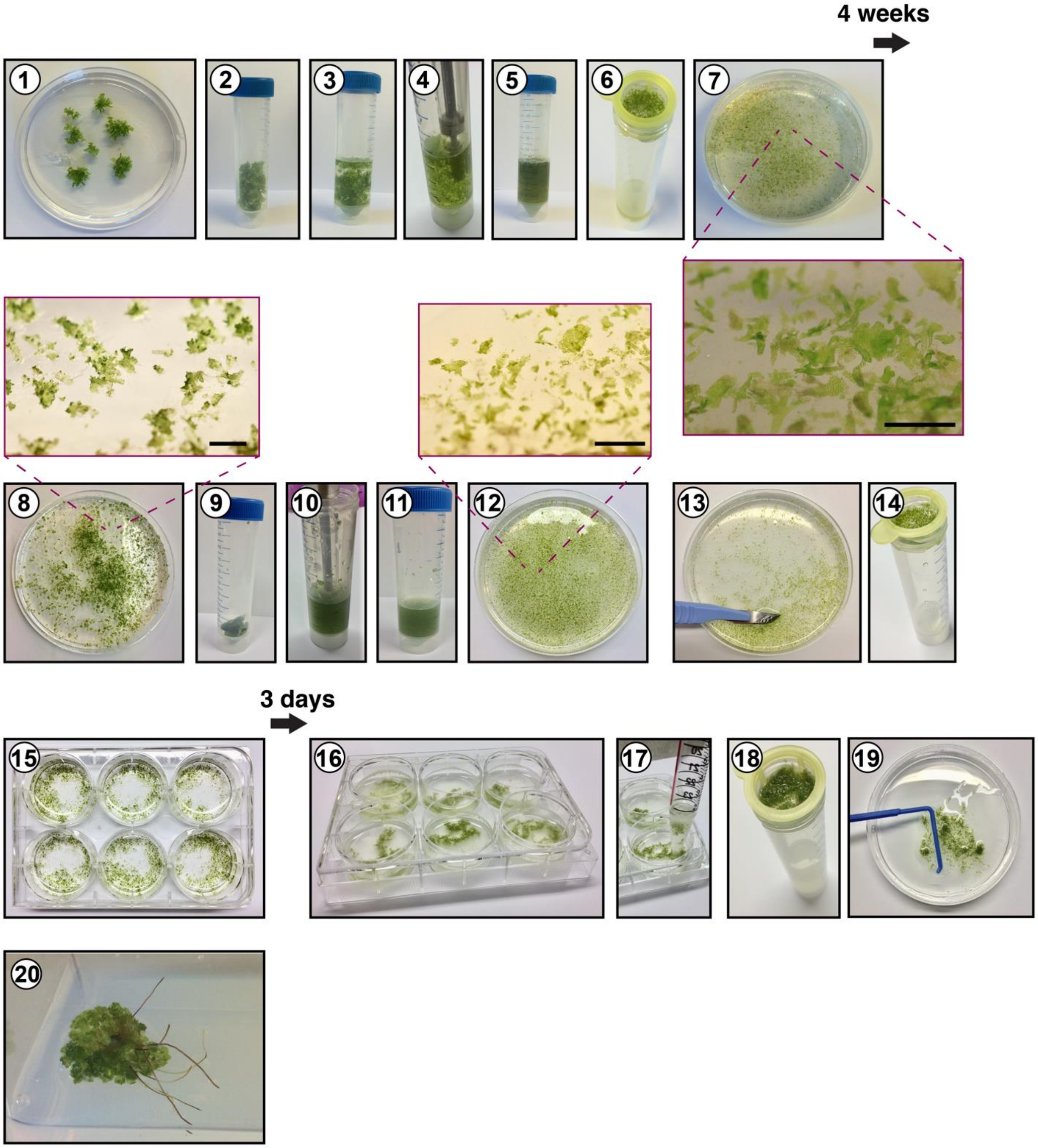
4 weeks
1
2
3
4
5
6
7
8
9
10
11
12
13
14
3 days
15
16
17
18
19
20

**Supplemental Figure S7: *A. agrestis* Bonn transformation protocol**

**1)** Approximately 2 g of thallus tissue grown for 3-4 weeks under low light intensity (similar to Supplemental Fig. S1B and Supplemental Fig. S2B) were collected. **2-5)** Tissue was split into three parts, transferred to three 50 mL falcon tubes containing 15 mL of sterile water and homogenized using a homogenizer and corresponding dispensing tools (for 5 sec, lowest power, 8000rpm). **6)** The tissue was washed using a cell strainer and **7)** transferred onto solid KNOP medium for further cultivation at 21°C under 12 hours light and 12 hours dark, light intensity 3-5 μmol $m^{-2}$ $s^{-1}$. **8-9)** After 3 weeks the tissue was transferred into a falcon tube using sterile scalpels, **10-11)** re-homogenized in 15-20 mL of sterile water and filtered using a cell strainer **12)** moved again onto 4 plates with solid KNOP medium and was allowed to grow for 2 days at 21°C under continuous light (intensity 30 μmol $m^{-2}$ $s^{-1}$). 5 mL of 2 days *Agrobacterium* culture was centrifuged for 7 min at 1800 xg. **13)** The regenerating thallus tissue was transferred into a 6-well plate (transfer ½ the tissue from one plate into a single well) with 4 mL of liquid KNOP medium supplemented with 2% (w/v) sucrose. 80 μL of *Agrobacterium* culture and acetosyringone at final concentration of 100 μM were added to the medium. 14) The tissue was co-cultivated with the *Agrobacterium* for 3 days on a shaker at 110 rpm, with only ambient light (1-3 μmol $m^{-2}$ $s^{-1}$). **15-17)** Using a sterile plastic pipette the tissue from a single well was transferred into a cell strainer, drained and then transferred onto growth media containing the appropriate antibiotic. To facilitate spreading of the tissue, 2 mL of sterile water was added to the petri dish. Plates were placed in a growth chamber at 22°C 12 hours light and 12 hours dark at light intensity of 35 μmol $m^{-2}$ $s^{-1}$. (**18)** After 3 weeks, the tissue was transferred to fresh growth media containing appropriate antibiotics. To facilitate spreading of the tissue, 2 mL of sterile water was added to the petri dish. Plates were placed in a growth chamber at 22°C 12 hours light and 12 hours dark at light intensity of 35 μmol $m^{-2}$ $s^{-1}$. **19)** After 6-8 weeks successful transformants were visible on the plate. To eliminate false positive regeneration, surviving tissue fragments were transferred again on antibiotics containing growth media. **20)** Transgenic plant with sporophytes.

Petri dish dimensions: 92 x16 mm

Scale bars: 2 mm.

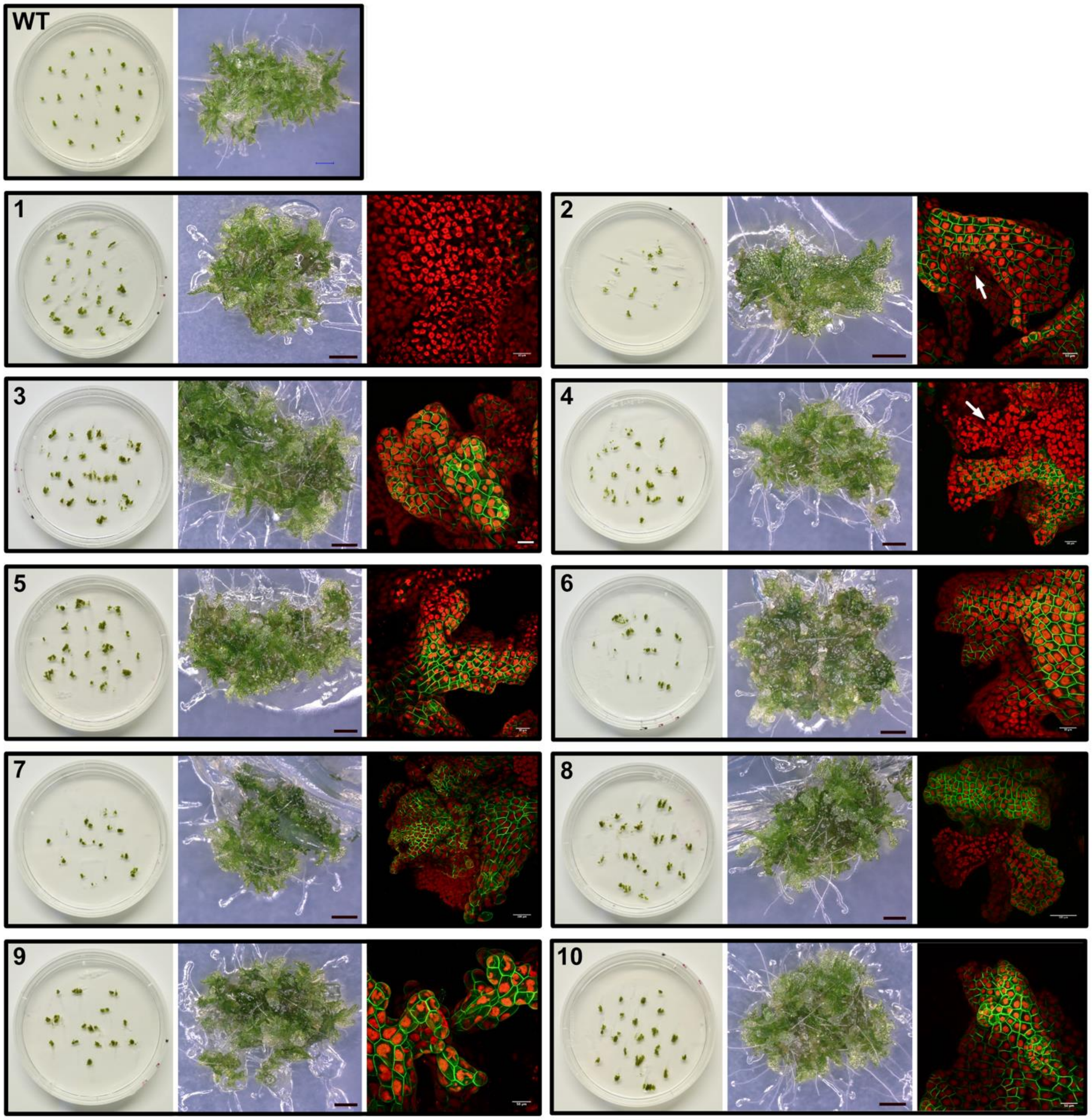
WT
1
2
3
4
5
6
7
8
9
10

**Supplemental Figure S8: Examples of *A. agrestis* Oxford transgenic lines expressing the *p-35S_s::eGFP-LTI6B* transcription unit**

Left: Examples of wild type and 10 independent transgenic *A. agrestis* lines (gametophyte thallus) transformed with the *p-35S::hph - p-35S_s::eGFP-LTI6B* construct. Petri dish dimensions: 92 x16 mm. Centre: Light micrograph of transgenic thallus. Scale bars: 500 μm. Right: Confocal microscopy images of thallus expressing the eGFP, driven by the CaMV 35S promoter, in the plasma membrane. Scale bars: 50 μm (7 and 8 scale bar: 100 μm). Younger parts of the thallus in 2 and 4 are indicated with white arrows.

### *AaEF1a*

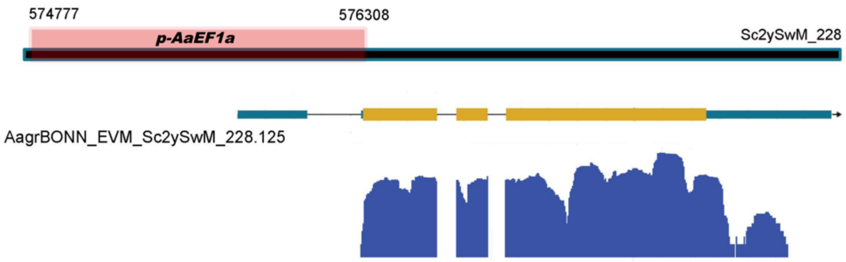


### *AaUbi*

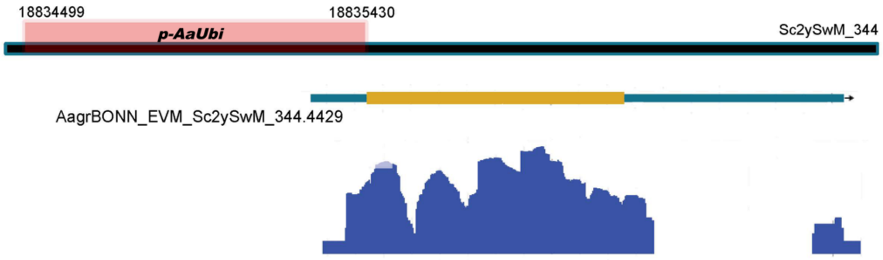


### *AaTip1;1*

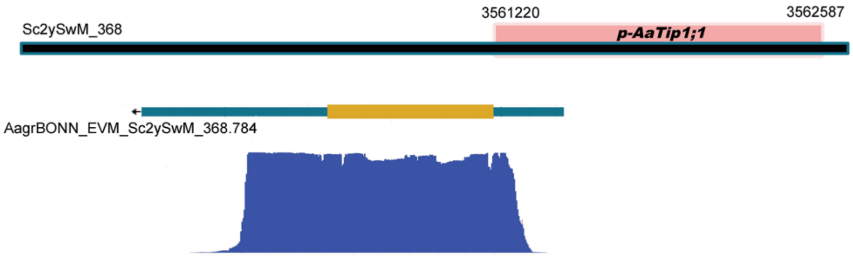


### *AaActin*

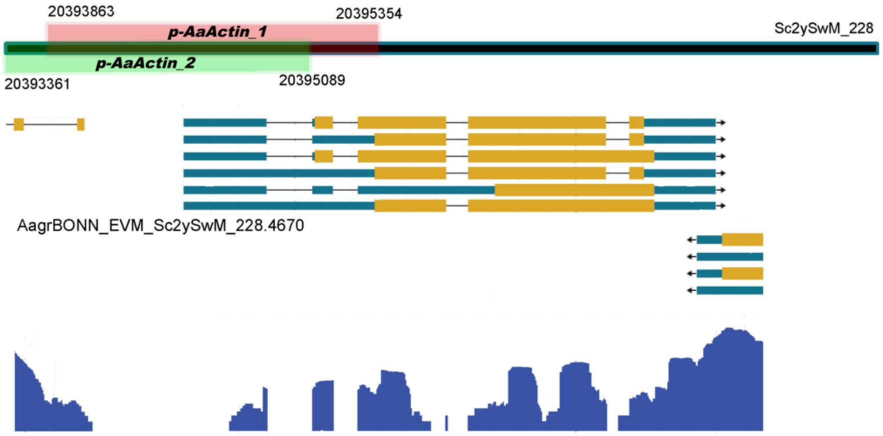


14

**Supplemental Figure S9: Promoters and their localization on the *A. agrestis* Bonn genome sequence**

Genomic scaffolds are shown as back lines at the top of each panel. Position of the promoter regions are represented with pink and green boxes (numbers refer to the end coordinates of the promoter sequences). Gene models are depicted below the genomic scaffolds (yellow box: exon, blue box: UTR, black line: intron). RNA-seq coverage track (log-scale) is visualized in blue at the bottom of each panel.

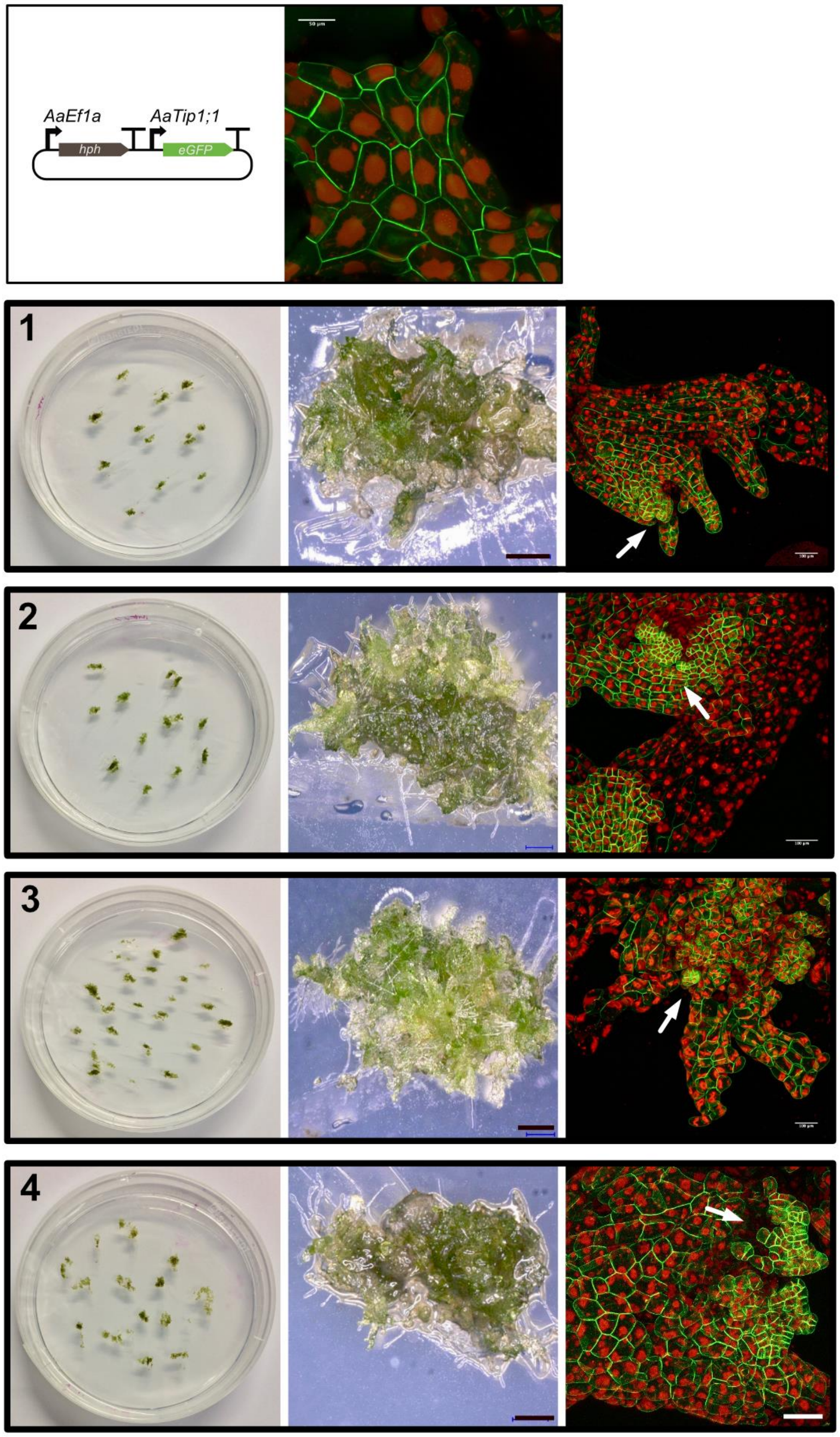
AaEf1a
AaTip1;1
hph
eGFP
1
2
3
4

**Supplemental Figure S10: Examples of *A. agrestis* Oxford transgenic lines expressing the *p-AaTip1;1::eGFP-LTI6B* or the *p-AaEF1a::eGFP-LTI6B* transcription unit**

Top: Example of an *A. agrestis* transgenic line (gametophyte thallus) transformed with the *p-AaEF1a::hph - p-AaTip1;1::eGFP-LTI6B* construct. Scale bar: 50 µm.

1-4 Left: Examples of 4 independent *A. agrestis* transgenic lines (gametophyte thallus) transformed with the *p-35S::hph - p-AaEF1a::eGFP-LTI6B* construct. Petri dish dimensions: 92 x16 mm. Centre: Light micrograph of transgenic thallus. Scale bar: 500 µm. Right: Confocal microscopy images of thallus expressing the eGFP, driven by the *AaEF1a* promoter, in the plasma membrane. Scale bars: 100 µm. Younger parts of the thallus are indicated with white arrows.

#### *A. agrestis* putative native promoter sequences

>*p-AaEF1a*

CACCGTCCAAACCTGGATAGCCGATATGTGAGTGTTTGGTTGCCTGTCATTTCTAAAAAGCGTATGTCTCGTAGC
GTAGATAGCAAGCGCTGTGAGAAAGGTGTGCTCCTATAGCATGGTAGCATCTGTCCCATATCTCGTCCGCGAGA
AGCCTACATTTCTCGTTGCGTATTAGAGCATGAAAAACCAACGTGAAGTACGTGATTCAGACCTTGACCAGTTCC
AGTTCTCTTCAACAAAAAAACAGTTCTTCAATTCATTTGCTTACATTAAACAATTGTCACACTCTTCTTCCCGGATG
ACAGCACCAAGACACTTCCCCTCCACAAAAATCTCCGAGTCATGTGGACCGGCCTGGATGCTCTTAAAGGGGTT
GCCATTGTAGCGGGGTGCCTTAACCACGACCTCTTCGATACCTGTCTGAAGCGGCGTGTGAAATCCTCCAGGAG
CTTGTTCAAATTTACAAGCTGCAATGGATGTAAACTCATCCAAAGACATGACTTGCGAGAAGCCAGAAACGGTCA
TCAACACTACGAACACCCCTAAACATGGATGGAGAGGCAACATCCTGCATGTGTCTATGTCAGAGGATGCAAAA
GTGAAAGACAAAGGAGAAGGATTGCTCGTGCAATCACTGCAAAGCAAGATGCGAATATCCCTAAAATCCACAGC
AGGAAAAGAAAGGAGCCTCAATAAGTCTCTAAAATGGGAAGAAAAGAAGCACGGGCCAGCGAGACGGAAATGA
GGCGATCCGATGCGATGGTTCCGAGAACTGAAGCGGTGCGTGGGGGGCTAGAAATGAATGGGGAACGTGGAG
GGCAAACAAAGGCTGTACTTAGCGGCAAATGCAGATGGCATGCATGCCAAGAACAGCAGCAGTCGGCACCGCC
AGTTTAGCGGAGAGCAACAGGCAGCAAAAGGAAGTGGAACAGGTTTAGGGCCGACGCAGTCGGCGCCTGCCCT
AAAACGGCAAGGCCCACGGCAGCGGTCGCGAGGGCGTTGCCTGCCGGTCGTGGAAAGCCGTGGGGCTGTCAA
CGGAACCTCACGCTCCCAGCCTGCCCTAAGCGGGGAAAAAGAAGAtCCACATTCTAGGGCTTTCCATCTCGGCT
CTCAGCCGTCCGATTAAGAATCGCACGGCCTAGCCCCGCCAGCCTAGCCCTCGGCCGCCCCCTTGACAGGTTA
AGAGCTGGGCAAGCGACTCAGTTGTTCCTTCGTCGCTCGCCCTTCTCACGAGCAGCTCGCCTGCCCCGCCGCC
GCCTTGCCTGCCTCGCGCCTTTTGTCTCCAGGTTCCCTTGCCGCGCCTTCTCCTTGGCTTGTGCTTCTGCTTTGC
TTGTGCCTCTGCTGGGCTCTACTTGTGTGGTGTTGTGCTTTGGCGCCGCCGCCTCGCTGCTGCTGTGGAGCTTG
GTGTGTGCTGGGGTTTCTGGGGTTTGCATGTGTCAGGGTTTGTAGATCTGTTCTTCGGGGCCGTCTTTCTAACGT
TTCGTGTGCCGTTGTTTCTGGGGGGTTTGGTGTGGAGCTCGCAGCTTGCAGCT

*>p-AaUbi*

TGACTGAGCAAGAATTCGTGCCTGGGAGTGAGCCAAGGCCTCTGATTGCAGAATATCGAACAAAGGAGGATTCT
CCACAAGCTGCAGAAGAAAAGTTTGATGTGTGAAGAAGAGAGGGAGAGAGAGACTTGCAGACAAATTCCATCAG
TCCTTAACCTAGTGTTGGCTGTATGATGGCTGACACCCGCTTCCTGTTATGCTCCTTTGCGTAGTGGAAAGCATA
CTCTGCAACGCGAGAGGTGGCCTGGCTGGTGATAATTTTTCAAGCTTTCCACACTGCGCTTCACTACCTGGACC
TGGCCAAAATCAGGTGACCGATCAGGCTGCAATGATAGCCACCATTCAAAATAAGCTGGCCAGTCTTCCAGCCT
CTCATCAGAGCTCGGGAACCACCGAAGTTGAATTGGGTTGAACAAGTGTCCCAACCCGATGCTCAAGGCCACCG
TACTCGCCTTCGGTTGTTTTCTCGGATGGTGTAAGATCATCCTCACATTTGCACAGAGCCGTAGCTTTTTCCGCT
GTCTCTGCTGGAAGAACGAATCAATTTGGCAACTGAGTTGGATGGTTCTTGTCAAACGGCTAGTTTGGCTGTTCA
GGCAAGGCCGATGGTGTTTTGCTATTTTAAAAGAGTTTTATTATCACCTACAAACCTACGTGGTAATCGAATATGT
GTGAGGTGGACAAGGAAAGTGGGCTGGAAGTTTCTGGAAAGAAGTATCCAGAACATCCATTCGCCGCTTTCCTG
GCTTGATTCACGCTTCGTCTTGCACGGCACTCCACCTCCTGGCCCAGTATAAATCCCTGCCTCCTTCCCTCTTTT
CTCACTCTCAGGGGTGTGAAGTTGTGGAGCGAGTCTGCTTGTCCTGAAGCTTGCTGCTAGTGGCGTCCTGCCCT
GCAATTACAGTCAGCTGTCCGTCTACCAAGTTTGCAAT

*>p-AaTip1;1*

CACCTGGAAGCATTTCCACACAGCATCTACACCACTACAGGGCTACATGCTCCATAGTGTAGAGCAAACCCTAGT
GCAGCTCAGATATTAGATAATGCTGTCCTTCCCAGAAAGCAAAAGGAATCGACAACTCTGGAAGGCAGACAGCA
CCAGTTCTGTGCAGCTTCATCCCGAGCTGATGCTGTCCTTCCGAGAAAGCAAAGGAATCGACACTCCATGGCCG
ACCGCACAAACACACCATAGATGCTGACCTATCTGGATATGCAACAACCACAACACTTGCGTCTCCACCGCTCCA
AGTCCAGCCTCCAGGAGCCAGGACTCACGGCCGGGTCGAGTGGGCCCAGCCTGGTCCCGTGGCGCCGGGGC
CCGGGCCCGGCTCCCCCCACTCCCCGGCGCGATTCTTACGCGGCGGACGGATAGGCCCACGACCGTCCTACG
TGGCGGGCCCCACGTAAGGACAGCGGTGCCTAGGCTGCACCAAATGCCGCGCCACCGTCCATGTCGTCAGAAT
CGGTCAGATATTAGCCCCTATCCCTTCACACGCCAGCGCCACGCCACGCCACGCTGCTGACGCCACCCCCCTC
TCACCATCATCATAGCCTTTCCAGATATCTCCACGCTCCTGCATTACCTGTACATCATTTTAATATGTGCATACATA
TGTATAATCAATCAGTAAAGGCAAGGCAGGCTGGATCCCCAGTAGAGCCCATCAATGCCTTGAATCTCGTGTTTC
CCGTCAGGTCCTTTCATTCATTCCAAGTCTCCTTCTCATACATACATCCCTCCGTGATTTAAACTTTATTAAATACC
TACATAGCAATGGAATAAATCTATAATGTGGAAGAGACTCCAAGGTGTGACTGAGGTTCCACATGTCTGAAAGAA
GGAGCATCAAAGTATCTACCTTCCCTTCCATTTGGTTGTATTCCCTTCATCCCCTAAACCCAATGCAAGTAGCAAA
AGAGGGACACGAGAAGGTAGTAGTGAAGGTATTGCATTATTGATTTGTGAAAGCGTTTGATGAAACACAAAGGG
CACACAAGAAGCGAATGAATGGATCCACACATGGAGGGGCAGGCAGCTCGAAAAACAGCAGAGGATGCACATG
GCCCAGAGCGGATAGGAAAGCAGGCCAGCCTCACTGGACAGTGGACAGCCAGGTGCAGCGCACCCTGGCATA
TTCGAACCGGAAGGGCTGACGTGGGCTCCATTATTTGGGGTGCTGAGTTGGCAAACACGCGCCGGGAAGCTCT
CAAAACCGGCATCTCCATCTCATTCCAACGCCGCAGTCCCATTGCTCACCAACCCCGGCTCCTTTCCCCGCTCT
GCTCCTCGGTTTCACGCTCCGCGCGAGCTCTGCTG

*>p-AaActin_1*

GTTCAGTCCTAACCGCGTTGTCAAAGCCCCACCCACTTGACCGCGGCCCGGTCAAGGTAACCGGGGCCCCCGC
GCTCCCATGTTACCGAATCTCGCCTCACCCCCGCTAGCCGGTTGCTTTCTTTAGGCCCTGGGGATGAAGCAAGG
CCGCGGACACGGGCCGGGGCCCACGAGCGAGTGGGTGTGGGCTGGACATGCCAGCCTTTTCTTCCTCTCAGT

GCTGCCTCCCTCACACCTGTGCCTTCTACGTGAAGTAACGTGAGTCATTGCCATTTTGTCGCTGACCAGGGTTTA
GGTTAGGGGAAACGAGTGTTGGAGTTAGGGCTTCTGCTCAACCACTGCACTGCTAGCGTGGGACGGCCGCGAT
TGCCCCGATGTTCCCTCCTGACGGCCACGATGCCTCTCCCGCCGCTGCGCGTACGGCCCCGCGCGTTCAGGC
GAATTCCGCTCGGCGAAACCCGCCCTAACCTAATCACTGCCCCACTCGCAAAACCGCAGGCCCGCCAACACGA
AGCCCGTCGTTTTTCGCACCAGAGGCTCAATTTTCCCCACACGCGTTTCGTGCGCGGCTTGTGCTTAACATTGC
CGCGCATTTATTGGCTGCACGATTCTTTTTCGAAAAATATCACGTTGGTTTCGCCAAGGTTTTCAAATGCAGGGG
CGGGTTTCGCCCGCGCATTCTCTCAGGCAGACGCAGTGCTCTCTCTCTCTTCCCCCGCAACTCGACATCACTGG
CTCCTCTCTTTCTCCGTTGTCTGCCCGCTTTCTCGCGCTCCTCTCCCGCTCGCCTCCGCGTTCCTGCACCACCG
CGGCTCGGCTCGCTCTCTCGCGCTTTATTGCCCGTGGGAGCGGGTTGCCTTGACAGTGTCGTGCCGCATCCAG
CAGCAGCGCGGAGCAGAGCAGAGCATCGCAAAGCACCGCAGAGCGGAGCCGGCCGGTGCGGAAGGCCGCAC
AGGCGAGGGCAAGGCGGTTGCTTGCGAGCGGCGCGGGCCCGATCGATCGTGTGCAGCAGGTGAGAGGGCCG
GTGGTGCCGGATCTGGCCTGGGAGGGAGCTGGATGGCTATCTTGTGTGTCTGTGTGGCGGTGGACTGATTTGC
AGGCTCGGATCTGGCTTTTTGCTGGTTCGATCTGAGTACATGAGCTTTCTTCCTGCATCTCGGCTGATCTTGGTT
CGCAATCTCGGGTTCCCGGCTGACGTGCATGTGGTGATCTGTAGGACTGAAAGAAATGGCCGAAAGATTGGCT
GGCGGTGGCGATCCCGAGGATATCCAGCCACTCGTCTGCGACAATGGTTCCGGAATGGTCAAGGTAAGAAGCT
ACCGCATCTAGATGTGGTATGGTATGGTTGTCCGGTTTGCATGTATGGAGGTGATTTTTAGGGTGCGGTTTTGGG
GAATTGGTTTGATGCGTTTTGCAGGCTGGATTTGCTGGAGATGATGCTCCCCGAGCTGTGTTCCCGAGTATCGT
GGGACGCCCGCGGCACACTGGTGTG

*>p-AaActin_2*

ACAGGGTTACTGCCTCTGGAGGCGTTCCTGGAAATCACAACGAGCAACCATTTTAGGTATCTCATCTTGCACTGT
TTTACAGTGAAGCATGTGTAAGCACAAGTAGCTGGTATTCAGCGCCAGTGACAGACAATCATCCCTCAATGTACT
CCCTCAGATCTGCTTCCTTAAAGCTCCATCAGCTGGTGCCAGGTGATGAAGCTTTGAAGACCGCAAACATATGA
GGATATTAACAACAAAACAAACAGGATGAAGAACATAGACAGCTTCCTCAAGCACTTGCGCGCTCGCAAGCAATC
CAAACCAACTTCACCTTCAGCCGTTCCACCTCAAGAATTCATGCTAAAGAGCAAAGCCGCCGCATCAACACCTCG
AGCAAGTCTGCCGGATCCCTTCCATCCGCAGTCTTCCGCGCCTAGCGCCGGCATCCTGACAAGCCAGCCGCTT
GGCCTCACAGTGCGGAACCGGTTCCGCAGTGGGAAAGCGAAACTGGATATTCCTGGTTCAGTCCTAACCGCGT
TGTCAAAGCCCCACCCACTTGACCGCGGCCCGGTCAAGGTAACCGGGGCCCCCGCGCTCCCATGTTACCGAAT
CTCGCCTCACCCCCGCTAGCCGGTTGCTTTCTTTAGGCCCTGGGGATGAAGCAAGGCCGCGGACACGGGCCG
GGGCCCACGAGCGAGTGGGTGTGGGCTGGACATGCCAGCCTTTTCTTCCTCTCAGTGCTGCCTCCCTCACACC
TGTGCCTTCTACGTGAAGTAACGTGAGTCATTGCCATTTTGTCGCTGACCAGGGTTTAGGTTAGGGGAAACGAGT
GTTGGAGTTAGGGCTTCTGCTCAACCACTGCACTGCTAGCGTGGGACGGCCGCGATTGCCCCGATGTTCCCTC
CTGACGGCCACGATGCCTCTCCCGCCGCTGCGCGTACGGCCCCGCGCGTTCAGGCGAATTCCGCTCGGCGAA
ACCCGCCCTAACCTAATCACTGCCCCACTCGCAAAACCGCAGGCCCGCCAACACGAAGCCCGTCGTTTTTCGCA
CCAGAGGCTCAATTTTCCCCACACGCGTTTCGTGCGCGGCTTGTGCTTAACATTGCCGCGCATTTATTGGCTGC
ACGATTCTTTTTCGAAAAATATCACGTTGGTTTCGCCAAGGTTTTCAAATGCAGGGGCGGGTTTCGCCCGCGCAT
TCTCTCAGGCAGACGCAGTGCTCTCTCTCTCTTCCCCCGCAACTCGACATCACTGGCTCCTCTCTTTCTCCGTTG
TCTGCCCGCTTTCTCGCGCTCCTCTCCCGCTCGCCTCCGCGTTCCTGCACCACCGCGGCTCGGCTCGCTCTCT
CGCGCTTTATTGCCCGTGGGAGCGGGTTGCCTTGACAGTGTCGTGCCGCATCCAGCAGCAGCGCGGAGCAGA
GCAGAGCATCGCAAAGCACCGCAGAGCGGAGCCGGCCGGTGCGGAAGGCCGCACAGGCGAGGGCAAGGCGG
TTGCTTGCGAGCGGCGCGGGCCCGATCGATCGTGTGCAGCAGGTGAGAGGGCCGGTGGTGCCGGATCTGGCC

TGGGAGGGAGCTGGATGGCTATCTTGTGTGTCTGTGTGGCGGTGGACTGATTTGCAGGCTCGGATCTGGCTTTT
TGCTGGTTCGATCTGAGTACATGAGCTTTCTTCCTGCATCTCGGCTGATCTTGGTTCGCAATCTCGGGTTCCCGG
CTGACGTGCATGTGGTGATCTGTAGGACTGAAAGAA

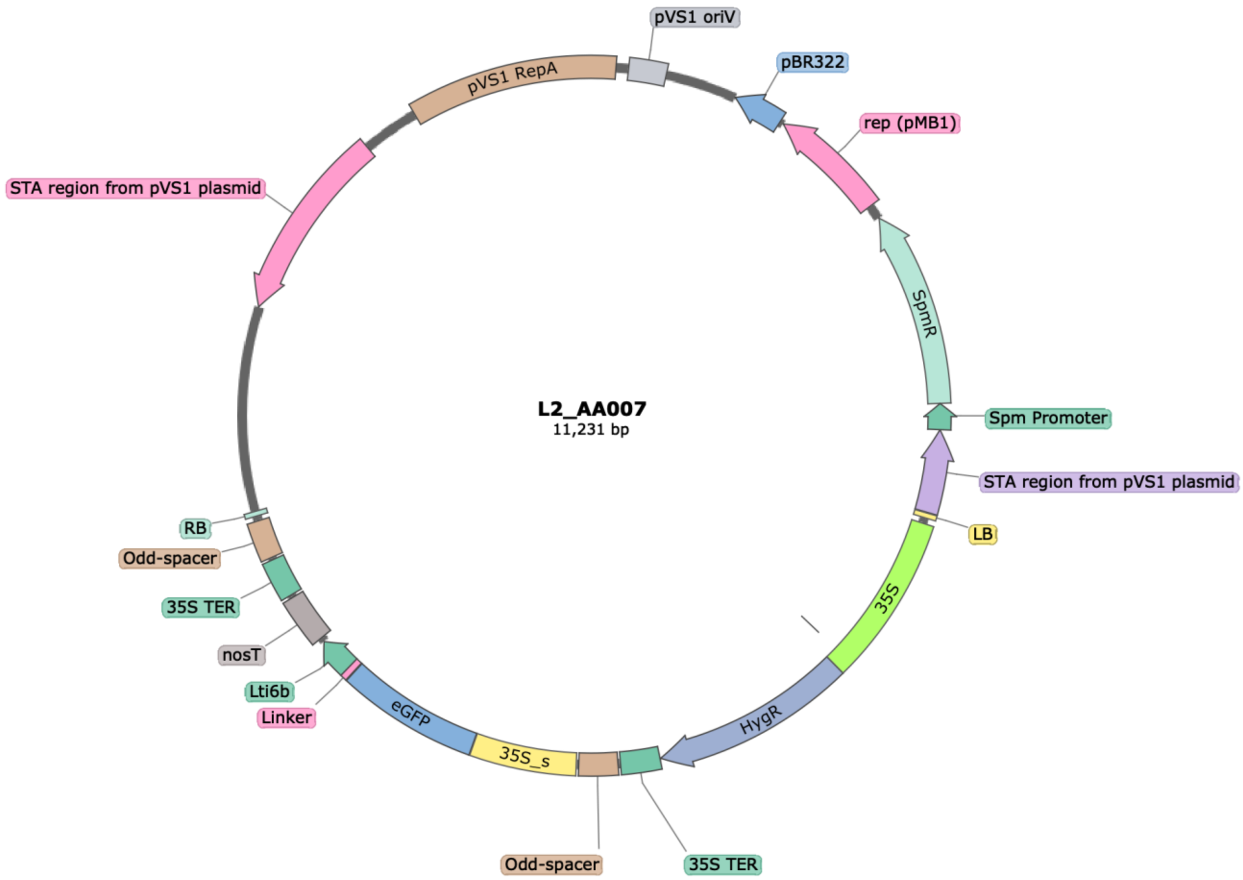


**Map of the transformation vector used for the generation of the *p-35S_s::eGFP-LTI6B* expressing lines** (created by SnapGene Viewer).

### >L2_AA007 sequence

tggcaggatatattgtggtgtaaacataacggatcCggtctcaggagaatgaggagAGATTAGCCTTTTCAATTTCAGAAAGAATGCTAACCCACAGATGG
TTAGAGAGGCTTACGCAGCAAGTCTCATCAAGACGATCTACCCGAGCAATAATCTACAGGAAATCAAATACCTTCCCAAGAAGG
TTAAAGATGCAGTCAAAAGATTCAGGACTAACTGCATCAAGAACACAGAGAAAGATATATTTCTCAAGATCAGAAGTACTATTCC
AGTATGGACGATTCAAGGCTTGCTTCACAAACCAAGGCAAGTAATAGAGATTGGAGTCTCTAAAAAGGTAGTTCCCACTGAATC
AAAGGCCATGGAGTCAAAGATTCAAATAGAGGACCTAACAGAACTCGCCGTAAAGACTGGCGAACAGTTCATACAGAGTCTCTT
ACGACTCAATGACAAGAAGAAAATCTTCGTCAACATGGTGGAGCACGACACACTTGTCTACTCCAAAAATATCAAAGATACAGTC
TCAGTAGACCAAAGGGCAATTGAGACTTTTCAACAAAGGGTAATATCCGGAAACCTCCTCGGATTCCATTGCCCAGCTATCTGT
CACTTTATTGTGAAGATAGTGGAAAAGGAAGGTGGCTCCTACAAATGCCATCATTGCGATAAAGGAAAGGCCATCGTTGAAGAT
GCCTCTGCCGACAGTGGTCCCAAAGATGGACCCCCACCCACGAGGAGCATCGTGGAAAAAGAAAACGTTCCAACCACGTCTAC
AAAGCAAGTGGATTGATGTGATATCTCCACTGACGTAAGGGATGACGCACAATCCCACTATCCTTCGCAAGACCCTTCCTCTAT
ATAAGGAAGTTCATTTCATTTGGAGAGAACACGGGGGACTCTAGAGGATCCAACATCACCAATGaaaaagcctgaactcaccgcgacgtctg
tcgagaagtttctgatcgaaaagttcgacagcgtctccgacctgatgcagctctcggagggcgaagaatctcgtgctttcagcttcgatgtaggagggcgtggatatgtcctgcgggtaaata
gctgcgccgatggtttctacaaagatcgttatgtttatcggcactttgcatcggccgcgctcccgattccggaagtgcttgacattggggagtttagcgagagcctgacctattgcatctcccgcc
gttcacagggtgtcacgttgcaagacctgcctgaaaccgaactgcccgctgttctacaaccggtcgcggaggctatggatgcgatcgctgcggccgatcttagccagacgagcgggttcg
gcccattcggaccgcaaggaatcggtcaatacactacatggcgtgatttcatatgcgcgattgctgatccccatgtgtatcactggcaaactgtgatggacgacaccgtcagtgcgtccgtcg
cgcaggctctcgatgagctgatgctttgggccgaggactgccccgaagtccggcacctcgtgcacgcggatttcggctccaacaatgtcctgacggacaatggccgcataacagcggtca
ttgactggagcgaggcgatgttcggggattcccaatacgaggtcgccaacatcttcttctggaggccgtggttggcttgtatggagcagcagacgcgctacttcgagcggaggcatccgga
gcttgcaggatcgccacgactccgggcgtatatgctccgcattggtcttgaccaactctatcagagcttggttgacggcaatttcgatgatgcagcttgggcgcagggtcgatgcgacgcaat
cgtccgatccggagccgggactgtcgggcgtacacaaatcgcccgcagaagcgcggccgtctggaccgatggctgtgtagaagtactcgccgatagtggaaaccgacgccccagcac
tcgtccgagggcaaagaaatagGCTTctctagctagagtcgatcgacaagctcgagtttctccataataatgtgtgagtagttcccagataagggaattagggttcctatagggtttcgctc
atgtgttgagcatataagaaacccttagtatgtatttgtatttgtaaaatacttctatcaataaaatttctaattcctaaaaCcaaaatccagtactaaaatccagatcgctagcaaggagTCT
GAGCAGGACAACTCGCGTAGTGAGAGTTACATGTTCGTTGGGTTCTTCCGACACGGACCTGAGTTGGCCAACGTCCCACCTGA
GGTCTGTGCCCCGGTGATGAGAAGTGTGCATCTCGTTCTTGCAGCTCGTCAGTACTTTCAGAATCATGGCGTGCATGGTAGAAT
GACCCTTATAACGGACTTCGACATGGCAATcgctatacaggagcatggagtcaaagattcaaatagaggacctaacagaactcgccgtaaagactggcgaacagtt
catacagagtctcttacgactcaatgacaagaagaaaatcttcgtcaacatggtggagcacgacacacttgtctactccaaaaatatcaaagatacagtctcagaagaccaaagggcaat
tgagacttttcaacaaagggtaatatccggaaacctcctcggattccattgcccagctatctgtcactttattgtgaagatagtggaaaaggaaggtggctcctacaaatgccatcattgcgat
aaaggaaaggccatcgttgaagatgcctctgccgacagtggtcccaaagatggacccccacccacgaggagcatcgtggaaaaagaagacgttccaaccacgtcttcaaagcaagt
ggattgatgtgatatctccactgacgtaagggatgacgcacaatcccactatccttcgcaagacccttcctctatataaggaagttcatttcatttggagagaacacgggggactAATGgtg
agcaagggcgaggagctgttcaccggggtggtgcccatcctggtcgagctggacggcgacgtaaacggccacaagttcagcgtgtccggcgagggcgagggcgatgccacctacgg
caagctgaccctgaagttcatctgcaccaccggcaagctgcccgtgccctggcccaccctcgtgaccaccctgacctacggcgtgcagtgcttcagccgctaccccgaccacatgaagc
agcacgacttcttcaagtccgccatgcccgaaggctacgtccaggagcgcaccatcttcttcaaggacgacggcaactacaagacccgcgccgaggtgaagttcgagggcgacaccct
ggtgaaccgcatcgagctgaagggcatcgacttcaaggaggacggcaacatcctggggcacaagctggagtacaactacaacagccacaacgtctatatcatggccgacaagcaga
agaacggcatcaaggtgaacttcaagatccgccacaacatcgaggacggcagcgtgcagctcgccgaccactaccagcagaacacccccatcggcgacggccccgtgctgctgccc
gacaaccactacctgagcacccagtccgccctgagcaaagaccccaacgagaagcgcgatcacatggtcctgctggagttcgtgaccgccgccgggatcactctcggcatggacgag
ctgtacaaggcttcgggagctgcggcagctgccgctgcggcagcggccgaattcaagcgcttgaaaatgagtacagccactttcgtagagattattcttgctatcatcttgcctcctctcggcg
tctttctcaaatttggttgcaaggttgagttttggatatgtttgattttgacgctgtttggttatcttcccggaatcctttacgctctttatatcatcaccttttgatgagcttgagctcgaatttccccgatcgtt
caaacatttggcaataaagtttcttaagattgaatcctgttgccggtcttgcgatgattatcatataatttctgttgaattacgttaagcatgtaataattaacatgtaatgcatgacgttatttatgaga
tgggtttttatgattagagtcccgcaattatacatttaatacgcgatagaaaacaaaatatagcgcgcaaactaggataaattatcgcgcgcggtgtcatctatgttactagatcGATCCGT
ATCGATAGCctctagctagagtcgatcgacaagctcgagtttctccataataatgtgtgagtagttcccagataagggaattagggttcctatagggtttcgctcatgtgttgagcatataa
gaaacccttagtatgtatttgtatttgtaaaatacttctatcaataaaatttctaattcctaaaaTcaaaatccagtactaaaatccagatCGCTacagaggagTCTGAGCAGGACA
ACTCGCGTAGTGAGAGTTACATGTTCGTTGGGTTCTTCCGACACGGACCTGAGTTGGCCAACGTCCCACCTGAGGTCTGTGCC
CCGGTGATGAGAAGTGTGCATCTCGTTCTTGCAGCTCGTCAGTACTTTCAGAATCATGGCGTGCATGGTAGAATGACCCTTATA
ACGGACTTCGACATGGCAATcgctaggtatacttgagaccGGATCCtgacaggatatattggcgggtaaac

### T-DNA

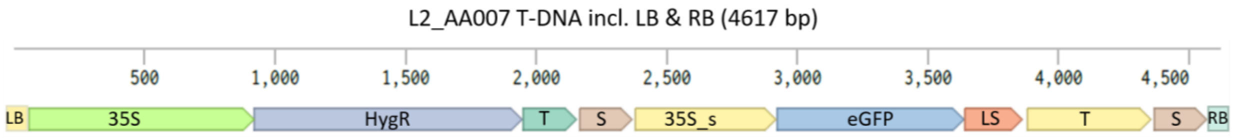


**Detailed description of insertion localization in the sequenced transformant lines**

***--Line Cam1:***

Based on our blastn search, we inferred that the insert is located on scaffold utg000058l (5`: 1418365 3’: 1418395, in the predicted 3’ -UTR of the *AagrOXF_EVM_utg000058l.283* locus) of the *A. agrestis* Oxford genome sequence (Li et al., 2020) [genome browser, blast interface and data are accessible under https://www.hornworts.uzh.ch/en/hornwort-genomes.html]. More specifically, we detected two inserts in head-to-head orientation, with one insert slightly truncated at the right border (missing 148 bp (part of 2$^{nd}$ spacer), the other one severely truncated at the right border (only the left border of the target sequence and part of the 35S-promoter are integrated).

***--Line Cam2:***

Our assembly showed that the T-DNA insert is located on scaffold utg000012l of the *A. agrestis* Oxford isolate’s genome (5’: 1410822 3’: 1410834). The insert is full length, but 12 bp are missing from the flanking genomic region. The insert is located immediately upstream of the predicted 5’ -UTR of the gene model *AagrOXF_EVM_utg000012l.314*.

***--Line BTI1:***

We found that the T-DNA insert is located on the scaffold utg000109l of the *A. agrestis* Oxford isolate’s genome (5’: 884855 3’: 884868 (13 bp missing from insertion site). More specifically, the insert occurs in the predicted 3’UTR of the gene model *AagrOXF_EVM_utg000109l.207* and immediately upstream of 5’UTR of gene model *AagrOXF_EVM_utg000109l.208.* The sequence data suggests at least two inserts in a head-to-head inverted arrangement. The T-DNA sequences that border the genomic flanking regions are on two different scaffolds of the assembly and both those scaffolds are broken shortly after the left border LB (at the beginning of p35S-BP). Therefore, our assembly provides no information on what is inserted in between the two detected partial inserts. Nevertheless, it is likely that there are multiple repeated and to various degrees truncated inserts in inverse and tandem directions present, suggested by the occurrence of various scaffolds with truncated inverse and tandemly repeated T-DNA inserts.

***--Line BTI2:***

We found that the T-DNA insert was located on scaffold utg00020l of the *A. agrestis* Oxford isolate's genome (5': 1209061 3': 1209065) at the beginning of the predicted 5'UTR of gene model *AagrOXF_EVM_utg000020l.241* and immediately downstream of the 3'UTR of gene model *AagrOXF_EVM_utg000020l.242*. Similar to the BTI1 line, there are at least two tandemly integrated inserts at the same locus in a head-to-head arrangement. The T-DNA bordering genomic flanking regions are on different scaffolds of the assembly with one of the scaffolds broken shortly after the left border, the right one containing only a short part of the p35S-BP next to the genomic flanking sequence. The sequence bridging these two partial inserts could not be assembled into a longer sequence. Again the occurrence of various scaffolds with truncated inverse and tandemly repeated T-DNA inserts suggests the presence of multiple repeated and to various degrees truncated inserts in inverse and tandem directions.

***--Line BTI3:***

We found the insert to be integrated on scaffold utg000012l of the *A. agrestis* Oxford isolate's genome (5': 1350616 3': 1350607), downstream of gene model *AagrOXF_EVM_utg000012l.300*. Similar to lines BTI1 and BTI2, there are at least two inserts integrated at the same locus in a head-to-head arrangement. More specifically, we found the T-DNA sequences that border the genomic flanking regions on two separate scaffolds of the transformed line's genome assembly and both those scaffolds were broken shortly after the left border (within the sequence of the p35S). Therefore it is impossible to say what exactly is in between. However, an additional scaffold contains parts of two head-to-head inverted T-DNA stretches. The first part breaks off in the middle of the second terminator and is fused to the middle of the fist spacer of the second part running in the inverse direction. This suggests that the line BTI3 indeed contains two head to head inverted inserts, with one being truncated at the second terminator and the other one truncated at the first spacer. Thus, BTI3 possibly contains two functional transcriptional units expressing the *hygromycin B phosphotransferase* gene but only one functional transcriptional unit expressing *eGFP*. Unlike BTI1 and BTI2, there are no other scaffolds present that would suggest additional partial inserts.

| Genome | Scaffold | Content | Remarks |
|---|---|---|---|
| BTI1 | NODE_27538_length_161_cov_50.892857 | 1-78 contain T-DNA 4515-4593 (end of spacer to RB) 85-161 contain T-DNA 3683-3760 (part of LS Lti6B) in reverse direction | indicating inverse partial inserts |
| | NODE_42621_length_149_cov_28.916667 | 1-77 contain T-DNA 97-196 (part of 35S) in reverse direction 86-149 contain T-DNA 14-191 (LB and part of 35S) | indicating inverse partial inserts |
| | NODE_41179_length_151_cov_51.594595 | 1-76 contain T-DNA 4516-4593 (end of spacer to RB) 77-151 contain T-DNA 3449-3524 (part of eGFP CDS) in reverse direction | indicating inverse partial inserts |
| | NODE_40940_length_152_cov_22.746667 | 1-76 contain T-DNA 4516-4592 (end of spacer to RB) 77-152 contain T-DNA 4326-4403 (end of terminator to start of spacer) in reverse direction | indicating inverse partial inserts |
| | NODE_21482_length_189_cov_21.767857 | 1-78 contain T-DNA 3916-3993 (part of terminator) in reverse direction<br>113-189 contain T-DNA 8-85 (LB and part of 35S) in reverse direction | indicating tandem partial inserts |

| | | | |
|---|---|---|---|
| BTI2 | NODE_15258_length_228_cov_20.377483 | 1-77 contain T-DNA 2294-2371 (or 4494-4571) (part of spacer) 150-228 contain T-DNA 4514-4593 (end of spacer to RB) in reverse direction | indicating inverse partial inserts<br><br>Note: No blast hit for seq of scaffold from 78-149 |
| | NODE_24906_length_170_cov_33.000000 | 1-77 contain T-DNA 4514-4590 (end of spacer to RB) 97-170 contain T-DNA 19-101 (LB and part of 35S) in same direction | indicating tandem inserts |
| | NODE_24845_length_170_cov_37.774194 | 1-77 contain T-DNA 4295-4372 (end of terminator to beginning of spacer)<br>94-170 contain T-DNA 1071-1148 (part of HygR CDS) in reverse direction | indicating inverse partial inserts |
| BTI3 | NODE_3513_length_2530_cov_42.732259 | 1-1160 contain T-DNA 2918-4078 (beginning of eGFP CDS to mid of terminator)<br>1161-2530 contain T-DNA 899-2270 (end of 35S to middle of first spacer) in reverse direction | indicating inverse partial inserts |

**Sanger sequencing results from PCR junction amplicons of Cam1, Cam2, BTI1, BTI2, and BTI3**

Green: Expected *A. agrestis* Oxford genomic DNA sequence
Yellow: Left border (underlined) to 35S promoter of insert sequence
Grey: HygR CDS of insert sequence
Lilac: 2nd terminator of insert sequence
Orange: Spacer to right border (underlined) of insert sequence
Dark blue: eGFP CDS of insert sequence

Pink: Lti6b of insert sequence
Blue: different 35S fragment fused sequence
Purple: random insertion sequence

>Cam1-5' preF (green = utg000058l.283 3'UTR locus as predicted using Illumina seq)
GTGAAGCCGTAAATGACGTGATGGTTCCCCTTGTTTATGTGGTTTGTTTATTGTTGGGTATGTGTGTGTGTGTGTGCGCCCTTGC
TAAGGCAGTGTTGAGCAAAGTGAAGGGGTTGTGGTCTCCGTTTTTTTTGGGGCCCACTAGGTTATGTGGCTTGTGGCACGGAAT
AAGGAGGTATATATTGTGGTGTAAACATAACGGATCCGGTCTCAGGAGAATGAGGAGAGATTAGCCTTTTCAATTTCAGAAAGA
ATGCTAACCCACAGATGGTTAGAGAGGCTTACGCAGCAAGTCTCATCAAGACGATCTACCCGAGCAATAATCTACAGGAAATCA
AATACCTTCCCAAGAAGGTTAAAGATGCAGTCAAAAGATTCAGGACTAACTGCATCAAGAACACAGAGAAAGATATATTTCTCAA
GATCAGAAGTACTATTCCAGTATGGACGATTCAAGGCTTGCTTCACAAACCAAGGCAAGTAATAGAGATTGGAGTCTCTAAAAA
GGTAGTTCCCACTGAATCAAAGGCCATGGAGTCAAAGATTCAAATAGAGGACCTAACAGAACTCGCCGTAAAGACTGGCGAAC
AGTTCATACAGAGTCTCTTACGACTCAATGACAAGAAGAAAATCTTCGTCAACATGGTGGAGCACGACACACTTGTCTACTCCAA
AAATATCAAAGATACAGTCTCAGTAGACCAAAGGGCAATTGAGACTTTTCAACAAAGGGTAATATCCGGAAACCTCCTCGGATTC
CATTGCCCAGCTATCTGTCACTTTATTGTGAAGATAGTGGAAAAGGAAGGTGGCTCCTACAAATGCCATCATTGCGATAAAGGA
AAGGCCATCGTTGAAGATGCCTCTGCCGACAGTGGTCCCAAAGATGGACCCCCACCCACGAGGAGCATCGTGGAAAAAGAAAA
CGTTCCAACCACGTCTACAAAGCAAGTGGATTGATGTGATATCTCCACTGACGTAAGGGATGACGCACAATCCCACTATCCTTC
GCAAGACCCTTCCTCTATATAAGGAAGTTCATTTCATTTGGAGAGAACACGGGGGACTCTAGAGGATCCAACATCACCAATGAA
AAAGCCTGAACTCACCGCGACGTCTGTCGAGAAGTTTCTGATCGAAAA

>Cam1-5' startHygRR (green = utg000058l.283 3'UTR locus as predicted using Illumina seq)
TTTTTTTGGGGCCCACTAGGTTATGTGGCTTGTGGCACGGAATAAGGAGGTATATATTGTGGTGTAAACATAACGGATCCGGTC
TCAGGAGAATGAGGAGAGATTAGCCTTTTCAATTTCAGAAAGAATGCTAACCCACAGATGGTTAGAGAGGCTTACGCAGCAAGT
CTCATCAAGACGATCTACCCGAGCAATAATCTACAGGAAATCAAATACCTTCCCAAGAAGGTTAAAGATGCAGTCAAAAGATTCA
GGACTAACTGCATCAAGAACACAGAGAAAGATATATTTCTCAAGATCAGAAGTACTATTCCAGTATGGACGATTCAAGGCTTGCT
TCACAAACCAAGGCAAGTAATAGAGATTGGAGTCTCTAAAAAGGTAGTTCCCACTGAATCAAAGGCCATGGAGTCAAAGATTCA
AATAGAGGACCTAACAGAACTCGCCGTAAAGACTGGCGAACAGTTCATACAGAGTCTCTTACGACTCAATGACAAGAAGAAAAT
CTTCGTCAACATGGTGGAGCACGACACACTTGTCTACTCCAAAAATATCAAAGATACAGTCTCAGTAGACCAAAGGGCAATTGA
GACTTTTCAACAAAGGGTAATATCCGGAAACCTCCTCGGATTCCATTGCCCAGCTATCTGTCACTTTATTGTGAAGATAGTGGAA
AAGGAAGGTGGCTCCTACAAATGCCATCATTGCGATAAAGGAAAGGCCATCGTTGAAGATGCCTCTGCCGACAGTGGTCCCAA
AGATGGACCCCCACCCACGAGGAGCATCGTGGAAAAAGAAAACGTTCCAACCACGTCTACAAAGCAAGTGGATTGATGTGATA
TCTCCACTGACGTAAGGGATGACGCACAATCCCACTATCCTTCGCAAGACCCTTCCTCTATATAAGGAAGTTCATTTCATTTGGA
GAGAACACGGGGGACTCTAGAGGATCCAACATCACCAATGAAAAAGCCTGAACTCACCGCGACGTCTGTCGAGAAGTTTCTGA
TCGAAAAGTTCGACAGCGTCTCCGACCTGATGCAGCTCTCGGAGGGCGAAGAATCTC

>Cam1-3' end 2nd termF (green = utg000058l.283 3'UTR locus as predicted using Illumina seq)
GGGTTTCGCTCATGTGTTGAGCATATAAGAAACCCTTAGTATGTATTTGTATTTGTAAAATACTTCTATCAATAAAATTTCTAATTC
CTAAAATCAAAATCCAGTACTAAAATCCAGATCGCTACAGAGGAGTCTGAGCAGGACAACTCGCGTAGTGAGAGTTACATGTTC
GTTGGGTTCTTCCGACACGGACCTGAGTTGGCCAACGTCAGTGGAGATATCACATCAATCCACTTGCTTTGTAGACGTGGTTGG
AACGTTTTCTTTTTCCACGATGCTCCTCGTGGGTGGGGGTCCATCTTTGGGACCACTGTCGGCAGAGGCATCTTCAACGATGGC
CTTTCCTTTATCGCAATGATGGCATTTGTAGGAGCCACCTTCCTTTTCCACTATCTTCACAATAAAGTGACAGATAGCTGGGCAA
TGGAATCCGAGGAGGTTTCCGGATATTACCCTTTGTTGAAAAGTCTCAATTGCCCTTTGGTCTACTGAGACTGTATCTTTGATAT
TTTTGGAGTAGACAAGTGTGTCGTGCTCCACCATGTTGACGAAGATTTTCTTCTTGTCATTGAGTCGTAAGAGACTCTGTATGAA
CTGTTCGCCAGTCTTTACGGCGAGTTCTGTTAGGTCCTCTATTTGAATCTTTGACTCCATGGCCTTTGATTCAGTGGGAACTACC
TTTTTAGAGACTCCAATCTCTATTACTTGCCTTGGTTTGTGAAGCAAGCCTTGAATCGTCCATACTGGAATAGTACTTCTGATCTT
GAGAAATATATCTTTCTCTGTGTTCTTGATGCAGTTAGTCCTGAATCTTTTGACTGCATCTTTAACCTTCTTGGGAAGGTATTTGA
TTTCCTGTAGATTATTGCTCGGGTAGATCGTCTTGATGAGACTTGCTGCGTAAGCCTCTCTAACCATCTGTGGGTTAGCATTCTT
TCTGAAATTGAAAAGGCTAATCTCTCCTCATTCTCCTGAGACCGGATCCGTTATGTTTACACCACAATATATCCAGGTAAGCCAC
TTTGCCATATTAGGCCCACTCACTGTACATGGATAGGAAG-
AGAGAAAAAAAGTGTTGGGTGCTGTAGACGCCTTGCAAGCAGGCGCGCTTACAGGTAACATAACCTAACCTAGGGACTTT

>Cam1-3' preR (green = utg000058l.283 3'UTR locus as predicted using Illumina seq)
AAGAAACCCTTAGTATGTATTTGTATTTGTAAAATACTTCTATCAATAAAATTTCTAATTCCTAAAATCAAAATCCAGTACTAAAATC
CAGATCGCTACAGAGGAGTCTGAGCAGGACAACTCGCGTAGTGAGAGTTACATGTTCGTTGGGTTCTTCCGACACGGACCTGA
GTTGGCCAACGTCAGTGGAGATATCACATCAATCCACTTGCTTTGTAGACGTGGTTGGAACGTTTTCTTTTTCCACGATGCTCCT
CGTGGGTGGGGGTCCATCTTTGGGACCACTGTCGGCAGAGGCATCTTCAACGATGGCCTTTCCTTTATCGCAATGATGGCATTT
GTAGGAGCCACCTTCCTTTTCCACTATCTTCACAATAAAGTGACAGATAGCTGGGCAATGGAATCCGAGGAGGTTTCCGGATAT
TACCCTTTGTTGAAAAGTCTCAATTGCCCTTTGGTCTACTGAGACTGTATCTTTGATATTTTTGGAGTAGACAAGTGTGTCGTGCT
CCACCATGTTGACGAAGATTTTCTTCTTGTCATTGAGTCGTAAGAGACTCTGTATGAACTGTTCGCCAGTCTTTACGGCGAGTTC

TGTTAGGTCCTCTATTTGAATCTTTGACTCCATGGCCTTTGATTCAGTGGGAACTACCTTTTTAGAGACTCCAATCTCTATTACTT
GCCTTGGTTTGTGAAGCAAGCCTTGAATCGTCCATACTGGAATAGTACTTCTGATCTTGAGAAATATATCTTTCTCTGTGTTCTTG
ATGCAGTTAGTCCTGAATCTTTTGACTGCATCTTTAACCTTCTTGGGAAGGTATTTGATTTCCTGTAGATTATTGCTCGGGTAGAT
CGTCTTGATGAGACTTGCTGCGTAAGCCTCTCTAACCATCTGTGGGTTAGCATTCTTTCTGAAATTGAAAAGGCTAATCTCTCCT
CATTCTCCTGAGACCGGATCCGTTATGTTTACACCACAATATATCCAGGTAAGCCACTTTGCCATATTAGGCCCACTCACTGTAC
ATGGATAGGAAGAAGAGAAAAAAAGTGTTGGGTGCTGTAGACGCCTTGCAAGCAGGCGCGCTTACAGGTAACATAACCTAACC
TAGGGACTTTATACTAGTGAAACTATCCAGAGTGAGCAGGTGACAAGT

>Cam2-5' preF (green = upstream of utg000012l.314 as predicted using Illumina seq)
GTCACTTCAGCTCTTGCTCTGGAGGGCAGTGAAATCTCTGGGAACTCGGTCCTCGTTCAAGAATGCGCCTGTCCGAGCACCAC
CAGCAACACAAGATTGAATACCACAGTTGCAACAAAAAAACCACACCATACATAGGTGGTGCTGCTCCTCTAGACTGTCACATC
CTCTCCGTCCGAGTAAATGCCTTAGGATATATTGTGGTGTAAACATAACGGATCCGGTCTCAGGAGAATGAGGAGAGATTAGCC
TTTTCAATTTCAGAAAGAATGCTAACCCACAGATGGTTAGAGAGGCTTACGCAGCAAGTCTCATCAAGACGATCTACCCGAGCA
ATAATCTACAGGAAATCAAATACCTTCCCAAGAAGGTTAAAGATGCAGTCAAAAGATTCAGGACTAACTGCATCAAGAACACAGA
GAAAGATATATTTCTCAAGATCAGAAGTACTATTCCAGTATGGACGATTCAAGGCTTGCTTCACAAACCAAGGCAAGTAATAGAG
ATTGGAGTCTCTAAAAAGGTAGTTCCCACTGAATCAAAGGCCATGGAGTCAAAGATTCAAATAGAGGACCTAACAGAACTCGCC
GTAAAGACTGGCGAACAGTTCATACAGAGTCTCTTACGACTCAATGACAAGAAGAAAATCTTCGTCAACATGGTGGAGCACGAC
ACACTTGTCTACTCCAAAAATATCAAAGATACAGTCTCAGTAGACCAAAGGGCAATTGAGACTTTTCAACAAAGGGTAATATCCG
GAAACCTCCTCGGATTCCATTGCCCAGCTATCTGTCACTTTATTGTGAAGATAGTGGAAAAGGAAGGTGGCTCCTACAAATGCC
ATCATTGCGATAAAGGAAAGGCCATCGTTGAAGATGCCTCTGCCGACAGTGGTCCCAAAGATGGACCCCCACCCACGAGGAGC
ATCGTGGAAAAAGAAAACGTTCCAACCACGTCTACAAAGCAAGTGGATTGATGTGATATCTCCACTGACGTAAGGGATGACGCA
CAATCCCACTATCCTTCGCAAGACCCTTCCTCTATATAAGGAAGTTCATTTCATTTGGAGAGAACACGGGGGACTCTAGAGGAT
CCAACATCACCAATGAAAAAGCCTGAA

>Cam2-5' startHygRR (green = upstream of utg000012l.314 as predicted using Illumina seq)
AAAAAACCACACCATACATAGGTGGTGCTGCTCCTCTAGACTGTCACATCCTCTCCGTCCGAGTAAATGCCTTAGGATATATTGT
GGTGTAAACATAACGGATCCGGTCTCAGGAGAATGAGGAGAGATTAGCCTTTTCAATTTCAGAAAGAATGCTAACCCACAGATG
GTTAGAGAGGCTTACGCAGCAAGTCTCATCAAGACGATCTACCCGAGCAATAATCTACAGGAAATCAAATACCTTCCCAAGAAG
GTTAAAGATGCAGTCAAAAGATTCAGGACTAACTGCATCAAGAACACAGAGAAAGATATATTTCTCAAGATCAGAAGTACTATTC
CAGTATGGACGATTCAAGGCTTGCTTCACAAACCAAGGCAAGTAATAGAGATTGGAGTCTCTAAAAAGGTAGTTCCCACTGAAT
CAAAGGCCATGGAGTCAAAGATTCAAATAGAGGACCTAACAGAACTCGCCGTAAAGACTGGCGAACAGTTCATACAGAGTCTCT
TACGACTCAATGACAAGAAGAAAATCTTCGTCAACATGGTGGAGCACGACACACTTGTCTACTCCAAAAATATCAAAGATACAGT
CTCAGTAGACCAAAGGGCAATTGAGACTTTTCAACAAAGGGTAATATCCGGAAACCTCCTCGGATTCCATTGCCCAGCTATCTG
TCACTTTATTGTGAAGATAGTGGAAAAGGAAGGTGGCTCCTACAAATGCCATCATTGCGATAAAGGAAAGGCCATCGTTGAAGA
TGCCTCTGCCGACAGTGGTCCCAAAGATGGACCCCCACCCACGAGGAGCATCGTGGAAAAAGAAAACGTTCCAACCACGTCTA
CAAAGCAAGTGGATTGATGTGATATCTCCACTGACGTAAGGGATGACGCACAATCCCACTATCCTTCGCAAGACCCTTCCTCTA
TATAAGGAAGTTCATTTCATTTGGAGAGAACACGGGGGACTCTAGAGGATCCAACATCACCAATGAAAAAGCCTGAACTCACCG
CGACGTCTGTCGAGAAGTTTCTGATCGAAAAGTTCGACAGCGTCTCCGACCTGATGCAGCTCTCGGAGGGCGAAGAATCTC

>Cam2-3' end eGFPF (green = upstream of utg000012l.314 as predicted using Illumina seq)
CGATCACATGGTCCTGCTGGAGTTCGTGACCGCCGCCGGGATCACTCTCGGCATGGACGAGCTGTACAAGGCTTCGGGAGCT
GCGGCAGCTGCCGCTGCGGCAGCGGCCGAATTCAAGCGCTTGAAAATGAGTACAGCCACTTTCGTAGAGATTATTCTTGCTAT
CATCTTGCCTCCTCTCGGCGTCTTTCTCAAATTTGGTTGCAAGGTTGAGTTTTGGATATGTTTGATTTTGACGCTGTTTGGTTATC
TTCCCGGAATCCTTTACGCTCTTTATATCATCACCTTTTGATGAGCTTGAGCTCGAATTTCCCCGATCGTTCAAACATTTGGCAAT
AAAGTTTCTTAAGATTGAATCCTGTTGCCGGTCTTGCGATGATTATCATATAATTTCTGTTGAATTACGTTAAGCATGTAATAATTA
ACATGTAATGCATGACGTTATTTATGAGATGGGTTTTTATGATTAGAGTCCCGCAATTATACATTTAATACGCGATAGAAAACAAA
ATATAGCGCGCAAACTAGGATAAATTATCGCGCGCGGTGTCATCTATGTTACTAGATCGATCCGTATCGATAGCCTCTAGCTAG
AGTCGATCGACAAGCTCGAGTTTCTCCATAATAATGTGTGAGTAGTTCCCAGATAAGGGAATTAGGGTTCCTATAGGGTTTCGC
TCATGTGTTGAGCATATAAGAAACCCTTAGTATGTATTTGTATTTGTAAAATACTTCTATCAATAAAATTTCTAATTCCTAAAATCAA
AATCCAGTACTAAAATCCAGATCGCTACAGAGGAGTCTGAGCAGGACAACTCGCGTAGTGAGAGTTACATGTTCGTTGGGTTCT
TCCGACACGGACCTGAGTTGGCCAACGTCCCACCTGAGGTCTGTGCCCCGGTGATGAGAAGTGTGCATCTCGTTCTTGCAGCT
CGTCAGTACTTTCAGAATCATGGCGTGCATGGTAGAATGACCCTTATAACGGACTTCGACATGGCAATCGCTAGGTATACTTGA
GACCGGATCCTGAAGCCGAGGATGAGTAAGCCGGGGATGTAGCTCAGATGGTAGAGCGCTCGTTTAGCATCCGAGGGAATCG
ATACCCGCATCTCCATGATTTGATTTGAAAGATGATAAAATAACTTTTCAACCGGAACGTC

>Cam2-3' preR (green = upstream of utg000012l.314 as predicted using Illumina seq)
CAGCCACTTTCGTAGAGATTATTCTTGCTATCATCTTGCCTCCTCTCGGCGTCTTTCTCAAATTTGGTTGCAAGGTTGAGTTTTG
GATATGTTTGATTTTGACGCTGTTTGGTTATCTTCCCGGAATCCTTTACGCTCTTTATATCATCACCTTTTGATGAGCTTGAGCTC
GAATTTCCCCGATCGTTCAAACATTTGGCAATAAAGTTTCTTAAGATTGAATCCTGTTGCCGGTCTTGCGATGATTATCATATAAT
TTCTGTTGAATTACGTTAAGCATGTAATAATTAACATGTAATGCATGACGTTATTTATGAGATGGGTTTTTATGATTAGAGTCCCG
CAATTATACATTTAATACGCGATAGAAAACAAAATATAGCGCGCAAACTAGGATAAATTATCGCGCGCGGTGTCATCTATGTTAC

TAGATCGATCCGTATCGATAGCCTCTAGCTAGAGTCGATCGACAAGCTCGAGTTTCTCCATAATAATGTGTGAGTAGTTCCCAG
ATAAGGGAATTAGGGTTCCTATAGGGTTTCGCTCATGTGTTGAGCATATAAGAAACCCTTAGTATGTATTTGTATTTGTAAAATAC
TTCTATCAATAAAATTTCTAATTCCTAAAATCAAAATCCAGTACTAAAATCCAGATCGCTACAGAGGAGTCTGAGCAGGACAACTC
GCGTAGTGAGAGTTACATGTTCGTTGGGTTCTTCCGACACGGACCTGAGTTGGCCAACGTCCCACCTGAGGTCTGTGCCCCGG
TGATGAGAAGTGTGCATCTCGTTCTTGCAGCTCGTCAGTACTTTCAGAATCATGGCGTGCATGGTAGAATGACCCTTATAACGG
ACTTCGACATGGCAATCGCTAGGTATACTTGAGACCGGATCCTGAAGCCGAGGATGAGTAAGCCGGGGATGTAGCTCAGATGG
TAGAGCGCTCGTTTAGCATCCGAGGGAATCGATACCCGCATCTCCATGATTTGATTTGAAAGATGATAAAATAACTTTTCAACCG
GAACGTCTTGACGAGTAAAGTAACTTTCTTCCAATATTTTTGACGAATAGAGTGATTCTTTTGACTGTAAATCTATAACTGAGCGC
TAGCAGTACTACACATCAC

>BTI1-5' preF (green = utg000109l.207 3'UTR locus as predicted using Illumina seq)
TCTCAACAATGGATATATTGTGGTGTAAACATAACGGATCCGGTCTCAGGAGAATGAGGAGAGATTAGCCTTTTCAATTTCAGAA
AGAATGCTAACCCACAGATGGTTAGAGAGGCTTACGCAGCAAGTCTCATCAAGACGATCTACCCGAGCAATAATCTACAGGAAA
TCAAATACCTTCCCAAGAAGGTTAAAGATGCAGTCAAAAGATTCAGGACTAACTGCATCAAGAACACAGAGAAAGATATATTTCT
CAAGATCAGAAGTACTATTCCAGTATGGACGATTCAAGGCTTGCTTCACAAACCAAGGCAAGTAATAGAGATTGGAGTCTCTAAA
AAGGTAGTTCCCACTGAATCAAAGGCCATGGAGTCAAAGATTCAAATAGAGGACCTAACAGAACTCGCCGTAAAGACTGGCGAA
CAGTTCATACAGAGTCTCTTACGACTCAATGACAAGAAGAAAATCTTCGTCAACATGGTGGAGCACGACACACTTGTCTACTCCA
AAAATATCAAAGATACAGTCTCAGTAGACCAAAGGGCAATTGAGACTTTTCAACAAAGGGTAATATCCGGAAACCTCCTCGGATT
CCATTGCCCAGCTATCTGTCACTTTATTGTGAAGATAGTGGAAAAGGAAGGTGGCTCCTACAAATGCCATCATTGCGATAAAGG
AAAGGCCATCGTTGAAGATGCCTCTGCCGACAGTGGTCCCAAAGATGGACCCCCACCCACGAGGAGCATCGTGGAAAAAGAAA
ACGTTCCAACCACGTCTACAAAGCAAGTGGATTGATGTGATATCTCCACTGACGTAAGGGATGACGCACAATCCCACTATCCTT
CGCAAGACCCTTCCTCTATATAAGGAAGTTCAT

>BTI1-5' mid35sR (green = utg000109l.207 3'UTR locus as predicted using Illumina seq)
CTCTTCTTCTTGCGCAGGTTGTGAATAATGACAGGCCAAATGCAGAGTTGATTGCATGTTCTGGACAACACGCCAACGTTGTCA
TTCCCTCGACTCCATTAGCCAAACCTCTCTGATTGCTCTCAACAATGGATATATTGTGGTGTAAACATAACGGATCCGGTCTCAG
GAGAATGAGGAGAGATTAGCCTTTTCAATTTCAGAAAGAATGCTAACCCACAGATGGTTAGAGAGGCTTACGCAGCAAGTCTCA
TCAAGACGATCTACCCGAGCAATAATCTACAGGAAATCAAATACCTTCCCAAGAAGGTTAAAGATGCAGTCAAAAGATTCAGGA
CTAACTGCATCAAGAACACAGAGAAAGATATATTTCTCAAGATCAGAAGTACTATTCCAGTATGGACGATTCAAGGCTTGCTTCA
CAAACCAAGGCAAGTAATAGAGATTGGAGTCTCTAAAAAGGTAGTTCCCACTGAATCAAAGGCCATGGAGTCAAAGATTCAAAT
AGAGGACCTAACAGAACTCGCCGTAAAGACTGGCGAACAGTTCATACAGAGTCTCTTACGACTCAATGACAAGAAGAAAATCTT
CGTCAACATGGTGGAGCACGACACACTTGTCTACTCCAAAAATATCAAAGATACAGTCTCAGTAGACCAAAGGGCAATTGAGAC
TTTTCAACAAAGGGTAATATCCGGAAACCTCCTCGGATTCCATTGCCCAGCTATCTGTCACTTTATTGTGAAGATAGTGGAAAAG
GAAGGTGGCTCCTACAAATG

>BTI1-3' mid35sF (green = utg000109l.207 3'UTR locus as predicted using Illumina seq)
TTTGTAGGAGCCACCTTCCTTTTCCACTATCTTCACAATAAAGTGACAGATAGCTGGGCAATGGAATCCGAGGAGGTTTCCGGA
TATTACCCTTTGTTGAAAAGTCTCAATTGCCCTTTGGTCTACTGAGACTGTATCTTTGATATTTTTGGAGTAGACAAGTGTGTCGT
GCTCCACCATGTTGACGAAGATTTTCTTCTTGTCATTGAGTCGTAAGAGACTCTGTATGAACTGTTCGCCAGTCTTTACGGCGAG
TTCTGTTAGGTCCTCTATTTGAATCTTTGACTCCATGGCCTTTGATTCAGTGGGAACTACCTTTTTAGAGACTCCAATCTCTATTA
CTTGCCTTGGTTTGTGAAGCAAGCCTTGAATCGTCCATACTGGAATAGTACTTCTGATCTTGAGAAATATATCTTTCTCTGTGTTC
TTGATGCAGTTAGTCCTGAATCTTTTGACTGCATCTTTAACCTTCTTGGGAAGGTATTTGATTTCCTGTAGATTATTGCTCGGGTA
GATCGTCTTGATGAGACTTGCTGCGTAAGCCTCTCTAACCATCTGTGGGTTAGCATTCTTTCTGAAATTGAAAAGGCTAATCTCT
CCTCATTCTCCTGAGACCGGATCCGTTATGTTTACACCACAATATATCCAGTCTGTAAATATGTGTTAATTAATAGACTCTGTACA
GCAAACGATTGTGTAGCTCCAGAAT

>BTI1-3' preR (green = utg000109l.207 3'UTR locus as predicted using Illumina seq)
CGTTTTCTTTTTCCACGATGCTCCTCGTGGGTGGGGGTCCATCTTTGGGACCACTGTCGGCAGAGGCATCTTCAACGATGGCCT
TTCCTTTATCGCAATGATGGCATTTGTAGGAGCCACCTTCCTTTTCCACTATCTTCACAATAAAGTGACAGATAGCTGGGCAATG
GAATCCGAGGAGGTTTCCGGATATTACCCTTTGTTGAAAAGTCTCAATTGCCCTTTGGTCTACTGAGACTGTATCTTTGATATTTT
TGGAGTAGACAAGTGTGTCGTGCTCCACCATGTTGACGAAGATTTTCTTCTTGTCATTGAGTCGTAAGAGACTCTGTATGAACTG
TTCGCCAGTCTTTACGGCGAGTTCTGTTAGGTCCTCTATTTGAATCTTTGACTCCATGGCCTTTGATTCAGTGGGAACTACCTTT
TTAGAGACTCCAATCTCTATTACTTGCCTTGGTTTGTGAAGCAAGCCTTGAATCGTCCATACTGGAATAGTACTTCTGATCTTGA
GAAATATATCTTTCTCTGTGTTCTTGATGCAGTTAGTCCTGAATCTTTTGACTGCATCTTTAACCTTCTTGGGAAGGTATTTGATTT
CCTGTAGATTATTGCTCGGGTAGATCGTCTTGATGAGACTTGCTGCGTAAGCCTCTCTAACCATCTGTGGGTTAGCATTCTTTCT
GAAATTGAAAAGGCTAATCTCTCCTCATTCTCCTGAGACCGGATCCGTTATGTTTACACCACAATATATCCAGTCTGTAA

>BTI2-5' preF (green = utg000020l.241 5'UTR locus as predicted using Illumina seq)
TTCCTCTGCTCTACATAACGGATCCGGTCTCAGGAGAATGAGGAGAGATTAGCCTTTTCAATTTCAGAAAGAATGCTAACCCACA
GATGGTTAGAGAGGCTTACGCAGCAAGTCTCATCAAGACGATCTACCCGAGCAATAATCTACAGGAAATCAAATACCTTCCCAA
GAAGGTTAAAGATGCAGTCAAAAGATTCAGGACTAACTGCATCAAGAACACAGAGAAAGATATATTTCTCAAGATCAGAAGTACT

ATTCCAGTATGGACGATTCAAGGCTTGCTTCACAAACCAAGGCAAGTAATAGAGATTGGAGTCTCTAAAAAGGTAGTTCCCACT
GAATCAAAGGCCATGGAGTCAAAGATTCAAATAGAGGACCTAACAGAACTCGCCGTAAAGACTGGCGAACAGTTCATACAGAGT
CTCTTACGACTCAATGACAAGAAGAAAATCTTCGTCAACATGGTGGAGCACGACACACTTGTCTACTCCAAAAATATCAAAGATA
CAGTCTCAGTAGACCAAAGGGCAATTGAGACTTTTCAACAAAGGGTAATATCCGGAAACCTCCTCGGATTCCATTGCCCAGCTA
TCTGTCACTTTATTGTGAAGATAGTGGAAAAGGAAGGTGGCTCCTACAAATGCCATCATTGCGATAAAGGAAAGGCCATCGTTG
AAGATGCCTCTGCCGACAGTGGTCCCAAAGATGGACCCCCACCCACGAGGAGCATCGTGGAAAAAGAAAACGTTCCAACCACG
TCTACAAAGCAAGTGGATTGATGTGATATCTCCACTGACGTAAGGGATGACGCACAATCCCACTATCCTTCGCAAGACCCTTCC

>BTI2-5' mid35sR (green = utg000020l.241 5'UTR locus as predicted using Illumina seq)
GGCGACAGCCTGGCCACTCACATTGCGGGCTTGCGGGCACAGCCGGTGTGGGACCCTCCGCCTGCGTCGCGGGGAAGGCTC
CACTCCACTCCTCCTCGACTCTCCCCGGCAACGCGGCCCGCGGGCAGCTCCCCGGCCTCCCCTTCTTTTCCTCTGCTCTACAT
AACGGATCCGGTCTCAGGAGAATGAGGAGAGATTAGCCTTTTCAATTTCAGAAAGAATGCTAACCCACAGATGGTTAGAGAGGC
TTACGCAGCAAGTCTCATCAAGACGATCTACCCGAGCAATAATCTACAGGAAATCAAATACCTTCCCAAGAAGGTTAAAGATGCA
GTCAAAAGATTCAGGACTAACTGCATCAAGAACACAGAGAAAGATATATTTCTCAAGATCAGAAGTACTATTCCAGTATGGACGA
TTCAAGGCTTGCTTCACAAACCAAGGCAAGTAATAGAGATTGGAGTCTCTAAAAAGGTAGTTCCCACTGAATCAAAGGCCATGG
AGTCAAAGATTCAAATAGAGGACCTAACAGAACTCGCCGTAAAGACTGGCGAACAGTTCATACAGAGTCTCTTACGACTCAATG
ACAAGAAGAAAATCTTCGTCAACATGGTGGAGCACGACACACTTGTCTACTCCAAAAATATCAAAGATACAGTCTCAGTAGACCA
AAGGGCAATTGAGACTTTTCAACAAAGGGTAATATCCGGAAACCTCCTCGGATTCCATTGCCCAGCTATCTGTCACTTTATTGTG
AAGATAGTGGAAAAGGAAGGTGGCTCCTACAAA

>BTI2-3' mid35sF (green = utg000020l.241 5'UTR locus as predicted using Illumina seq; blue = different 35S promoter fragment)
AGAGGCATCTTCAACGATGGCCTTTCCTTTATCGCAATGATGGCATTTGTAGGAGCCACCTTCCTTTTCCACTATCTTCACAATA
AAGTGACAGATAGCTGGGCAATGGAATCCGAGGAGGTTTCCGGATATTACCCTTTGTTGAAAAGTCTCAATTGCCCTTTGGTCT
TCTGAGACTGTATCTTTGATATTTTTGGAGTAGACAAGTGTGTCGTGCTCCACCATGTTGACGAAGATTTTCTTCTTGTCATTGAG
TCGTAAGAGACTCTGTATGACCTACACATAATTATAACCTGCCCTTCCAACCCGACACTCTCTTCCTGCGGAGTCGATCAGCTCA
CACAGCGCTAAAAAGCTGGCTTTTCGTGTGAAGCACCTCATGAAATCTTCAAGCTCCCTCACATGGTAAGCATTCATCATAAAGC
TCCACAGAGATTGAGTACAACACTGTAACGTTCATCCTT

>BTI2-3' preR (green = utg000020l.241 5'UTR locus as predicted using Illumina seq; blue = different 35S promoter fragment)
TCCAAATGAAATGAACTTCCTTATATAGAGGAAGGGTCTTGCGAAGGATAGTGGGATTGTGCGTCATCCCTTACGTCAGTGGAG
ATATCACATCAATCCACTTGCTTTGTAGACGTGGTTGGAACGTTTTCTTTTTCCACGATGCTCCTCGTGGGTGGGGGTCCATCTT
TGGGACCACTGTCGGCAGAGGCATCTTCAACGATGGCCTTTCCTTTATCGCAATGATGGCATTTGTAGGAGCCACCTTCCTTTT
CCACTATCTTCACAATAAAGTGACAGATAGCTGGGCAATGGAATCCGAGGAGGTTTCCGGATATTACCCTTTGTTGAAAAGTCT
CAATTGCCCTTTGGTCTTCTGAGACTGTATCTTTGATATTTTTGGAGTAGACAAGTGTGTCGTGCTCCACCATGTTGACGAAGAT
TTTCTTCTTGTCATTGAGTCGTAAGAGACTCTGTATGACCTACACATAATTATAACCTGCCCTTCCAACCCGACACTCTCTTCCTG
CGGAGTCGATCAGCTCACACAGCGCTAAAAAGCTGGCTTTTCGTGTGAAGCACCT

>BTI3-5' preF (green = upstream of utg000012l.300 locus as predicted using Illumina seq; red = random insertion)
GCTATGTAGACTGAACAATTTGAAATGAAATGTACATGTATATGTTCATTTCATGAACATGTATTTCAAATTGTAAATTATATATTG
TGGTGTAAACATAACGGATCCGGTCTCAGGAGAATGAGGAGAGATTAGCCTTTTCAATTTCAGAAAGAATGCTAACCCACAGAT
GGTTAGAGAGGCTTACGCAGCAAGTCTCATCAAGACGATCTACCCGAGCAATAATCTACAGGAAATCAAATACCTTCCCAAGAA
GGTTAAAGATGCAGTCAAAAGATTCAGGACTAACTGCATCAAGAACACAGAGAAAGATATATTTCTCAAGATCAGAAGTACTATT
CCAGTATGGACGATTCAAGGCTTGCTTCACAAACCAAGGCAAGTAATAGAGATTGGAGTCTCTAAAAAGGTAGTTCCCACTGAA
TCAAAGGCCATGGAGTCAAAGATTCAAATAGAGGACCTAACAGAACTCGCCGTAAAGACTGGCGAACAGTTCATACAGAGTCTC
TTACGACTCAATGACAAGAAGAAAATCTTCGTCAACATGGTGGAGCACGACACACTTGTCTACTCCAAAAATATCAAAGATACAG
TCTCAGTAGACCAAAGGGCAATTGAGACTTTTCAACAAAGGGTAATATCCGGAAACCTCCTCGGATTCCATTGCCCAGCTATCT
GTCACTTTATTGTGAAGATAGTGGAAAAGGAAGGTGGCTCCTACAAATGCCATCATTGCGATAAAGGAAAGGCCATCGTTGAAG
ATGCCTCTGCCGACAGTGGTCCCAAAGATGGACCCCCACCCACGAGGAGCATCGTGGAAAAAGAAAACG

>BTI3-5' mid35sR (green = upstream of utg000012l.300 locus as predicted using Illumina seq; red = random insertion)
AAAGGAATAAAGAGTTAAATTTGACGAGGGTTCTCAAATTGCGCAGTCTACATAGAACCGGGGCTATGTAGACTGAACAATTTG
AAATGAAATGTACATGTATATGTTCATTTCATGAACATGTATTTCAAATTGTAAATTATATATTGTGGTGTAAACATAACGGATCCG
GTCTCAGGAGAATGAGGAGAGATTAGCCTTTTCAATTTCAGAAAGAATGCTAACCCACAGATGGTTAGAGAGGCTTACGCAGCA
AGTCTCATCAAGACGATCTACCCGAGCAATAATCTACAGGAAATCAAATACCTTCCCAAGAAGGTTAAAGATGCAGTCAAAAGAT
TCAGGACTAACTGCATCAAGAACACAGAGAAAGATATATTTCTCAAGATCAGAAGTACTATTCCAGTATGGACGATTCAAGGCTT
GCTTCACAAACCAAGGCAAGTAATAGAGATTGGAGTCTCTAAAAAGGTAGTTCCCACTGAATCAAAGGCCATGGAGTCAAAGAT
TCAAATAGAGGACCTAACAGAACTCGCCGTAAAGACTGGCGAACAGTTCATACAGAGTCTCTTACGACTCAATGACAAGAAGAA
AATCTTCGTCAACATGGTGGAGCACGACACACTTGTCTACTCCAAAAATATCAAAGATACAGTCTCAGTAGACCAAAGGGCAATT
GAGACTTTTCAACAAAGGGTAATATCCGGAAACCTCCTCGGATTCCATTGCCCAGCTATCTGTCACTTTATTGTGAAGATAGTGG
AAAAGGAAGGTGGCTCCTACAAATGCCATCATTGCGATAAAGGAAAGGCCATCGTTGAAGATGCCTCTGC

>BTI3-3’ mid35sF (green = upstream of utg000012I.300 locus as predicted using Illumina seq; red = random insertion)
GCATTTGTAGGAGCCACCTTCCTTTTCCACTATCTTCACAATAAAGTGACAGATAGCTGGGCAATGGAATCCGAGGAGGTTTCC
GGATATTACCCTTTGTTGAAAAGTCTCAATTGCCCTTTGGTCTACTGAGACTGTATCTTTGATATTTTTGGAGTAGACAAGTGTGT
CGTGCTCCACCATGTTGACGAAGATTTTCTTCTTGTCATTGAGTCGTAAGAGACTCTGTATGAACTGTTCGCCAGTCTTTACGGC
GAGTTCTGTTAGGTCCTCTATTTGAATCTTTGACTCCATGGCCTTTGATTCAGTGGGAACTACCTTTTTAGAGACTCCAATCTCTA
TTACTTGCCTTGGTTTGTGAAGCAAGCCTTGAATCGTCCATACTGGAATAGTACTTCTGATCTTGAGAAATATATCTTTCTCTGTG
TTCTTGATGCAGTTAGTCCTGAATCTTTTGACTGCATCTTTAACCTTCTTGGGAAGGTATTTGATTTCCTGTAGATTATTGCTCGG
GTAGATCGTCTTGATGAGACTTGCTGCGTAAGCCTCTCTAACCATCTGTGGGTTAGCATTCTTTCTGAAATTGAAAAGGCTAATC
TCTCCTCATTCTCCTGAGACCGGATCCGTTATATGAAAACATACATGAAATTTGTACAATTTGAAAACTAATTTTCTGGACCCTTA
TCACGGCCTGCAGTTTTCGTTCTGGTTTATTCGCATATCGACCTCTCAGTTGGGACATGTGAACCGATGACTTGGGTGCTCCAG
TTCCCGGAGCTGCTGCATGTTA

>BTI3-3’ preR (green = upstream of utg000012I.300 locus as predicted using Illumina seq; red = random insertion)
CCACGATGCTCCTCGTGGGTGGGGGTCCATCTTTGGGACCACTGTCGGCAGAGGCATCTTCAACGATGGCCTTTCCTTTATCG
CAATGATGGCATTTGTAGGAGCCACCTTCCTTTTCCACTATCTTCACAATAAAGTGACAGATAGCTGGGCAATGGAATCCGAGG
AGGTTTCCGGATATTACCCTTTGTTGAAAAGTCTCAATTGCCCTTTGGTCTACTGAGACTGTATCTTTGATATTTTTGGAGTAGAC
AAGTGTGTCGTGCTCCACCATGTTGACGAAGATTTTCTTCTTGTCATTGAGTCGTAAGAGACTCTGTATGAACTGTTCGCCAGTC
TTTACGGCGAGTTCTGTTAGGTCCTCTATTTGAATCTTTGACTCCATGGCCTTTGATTCAGTGGGAACTACCTTTTTAGAGACTC
CAATCTCTATTACTTGCCTTGGTTTGTGAAGCAAGCCTTGAATCGTCCATACTGGAATAGTACTTCTGATCTTGAGAAATATATCT
TTCTCTGTGTTCTTGATGCAGTTAGTCCTGAATCTTTTGACTGCATCTTTAACCTTCTTGGGAAGGTATTTGATTTCCTGTAGATT
ATTGCTCGGGTAGATCGTCTTGATGAGACTTGCTGCGTAAGCCTCTCTAACCATCTGTGGGTTAGCATTCTTTCTGAAATTGAAA
AGGCTAATCTCTCCTCATTCTCCTGAGACCGGATCCGTTATATGAAAACATACATGAAATTTGTACAATTTGAAAACTAATTTTCT
G

**Supplemental Table I***:* List of lines generated

| | | | |
|---|---|---|---|
| ***Construct*** | *Total number of lines* | *Number of lines exhibiting fluorescence /GUS staining* | *Number of lines not exhibiting fluorescence /GUS staining* |
| pCAMBIA1305.1 | *10* | *10* | - |
| *p-35S::hph - p-35S_s::eGFP-LTI6b* | *157* | *153* | *4* |
| *p-35S::hph - p-35S_s::eGFP-LTI6b (*Bonn*)* | *2* | *2* | - |
| *p-35S::hph - p-35S_s::mTurquoise2-N7* | *6* | *5* | *1* |
| *p-35S::hph - p-35S_s::eYFP-myr* | *5* | *5* | - |
| *p-35S::hph - p-35S::mVenus-N7* | *3* | *3* | - |
| *p-35S::hph - p-AaEF1a::eGFP-LTI6b* | *8* | *6* | *2* |
| *p-35S::hph - p-AaUbi::eGFP-LTI6b* | *3* | *3* | - |
| *p-35S::hph - p-AaTip1;1::mTurquoise2-N7* | *3* | *3* | - |
| *p-EF1a::hph- p-AaTip1;1::eGFP-LTI6b* | *1* | *1* | - |
| *p-35S::hph - p-EF1a:mTurquoise2-N7 p-35S_s::eGFP-LTI6b* | *3* | *2* | *1* |

| | | | |
|---|---|---|---|
| *p-EF1a::hph- p-35S_s::eGFP-LTI6b* | *3* | *3* | - |
| *p-35S::hph - p-AaAct8_1::eGFP-LTI6b* | *3* | - | *3* |
| *p-35S::hph - p-AaAct8_2::eGFP-LTI6b* | *9* | - | *9* |

**Supplemental Table II***:*

| **Oxford** | |
|---|---|
| Gene | Gene ID |
| *EF1a* | AagrOXF_EVM_utg000056l.73 |
| *Tip1;1* | AagrOXF_EVM_utg000017l.183 |
| *Ubiquitin* (*Ubi*) | AagrOXF_EVM_utg000038l.265 |
| *Actin* | AagrOXF_EVM_utg000050l.84 |
| CN-Max 1 (N-plus1)* | AagrOXF_EVM_utg000083l.42 |
| CN-Max 2** | AagrOXF_EVM_utg000017l.207 |
| CN-Max 3 (N-plus2)* | AagrOXF_EVM_utg000095l.110 |
| CN-Max 4** | AagrOXF_EVM_.utg000024l.154 |
| CN-Max 5 (N-plus3)* | AagrOXF_EVM_utg000077l.83 |
| **Bonn** | |
| Gene | Gene ID |
| *EF1a* | AagrBONN_EVM_Sc2ySwM_228.125 |
| *Tip1;1* | AagrBONN_EVM_Sc2ySwM_368.784 |
| *Ubiquitin* (*Ubi*) | AagrBONN_EVM_Sc2ySwM_344.4429 |
| *Actin* | AagrBONN_EVM_Sc2ySwM_228.4670 |
| T-Max 1*** | AagrBONN_EVM_Sc2ySwM_368.2612 |
| T-Max 2*** | AagrBONN_EVM_Sc2ySwM_117.1320 |
| T-Max 3*** | AagrBONN_EVM_Sc2ySwM_344.2836 |
| T-Max 4*** | AagrBONN_EVM_Sc2ySwM_228.5295 |
| T-Max 5*** | AagrBONN_EVM_Sc2ySwM_344.2021 |

* Highest expressing genes based on N_plus conditions for Oxford

**Highest expressing genes based on C_normal conditions for Oxford

***Highest expressing genes under all conditions (across various developmental stages of the gametophyte and the sporophyte phases: young, intermediate and mature) for Bonn

| gene | Remark |
|---|---|
| CN-Max 1 | chlorophyll a/b-binding protein |
| CN-Max 2 | late embryogenesis abundant protein-related |
| CN-Max 3 | chlorophyll a/b-binding protein |
| CN-Max 4 | unknown |
| CN-Max 5 | unknown |
| T-Max 1 | LCIB (low-CO2 inducible protein LCIB [Chlamydomonas reinhardtii]) |
| T-Max 2 | chlorophyll a/b-binding protein (identical to CN-Max 3 Ox AagrOXF_evm.TU.utg000095l.110) |
| T-Max 3 | rbcS |
| T-Max 4 | Leucine-rich receptor-like protein kinase family protein |
| T-Max 5 | HSP20-like chaperones superfamily protein |

**Supplemental Table III***:* List of construct sequences (separate excel file)

**Supplemental Table IV***:* List of primers used in this study

| *primers* | | | | |
|---|---|---|---|---|
| ***1*** | ***AaEf1a*** | | ***AaEf1a-Domestication*** | |
| | *AA032-EF1A1-F1* | *caccgtccaaacctggatagc* | *AA047-EF1a-l-F1* | *gcgagctcttcgtctctggagcaccgtccaaacctggatagc* |
| | *AA035-EF1A1-R2* | *cctcttgttcatctcagcggc* | *AA052-EF1a-l-R3* | *tctagctcttcgtctcacattagctgcaagctgcgagctccac* |
| ***2*** | ***AaActin_1*** | | ***AaActin_1-Domestication*** | |
| | *AA027-actin8-F2* | *gttcagtcctaaccgcgttg* | *AA045-Act-l-F* | *gcgagctcttcgtctctggaggttcagtcctaaccgcgttgtc* |
| | *AA001-pro_actinR1* | *atccttctgtcccatgccaa* | *AA046-Act-l-R* | *tctagctcttcgtctcacattcacaccagtgtgccgcgggcg* |
| ***3*** | ***AaActin_2*** | | ***AaActin_2-Domestication*** | |
| | ***MW61_pAaAct1-F1*** | *agatccatccttccatcagcc* | ***MW63_pAaAct1-F2*** | *gtgagctcttcgtctctggaggaacagggttactgcctctgg* |
| | ***MW62_pAaAct1-R1*** | *cagacgagtggctggatatcc* | ***MW64_pAaAct1_R2*** | *gctagctcttcgtctcacatttctttcagtcctacagatcacc* |
| ***4*** | ***AaTip1;1*** | | | |
| | ***MW69_pAaTIP1-F1*** | *cgtccagcaccagcttactgg* | ***MW71-pAaTIP1-F2*** | *gtgagctcttcgtctctggagcacctggaagcatttccacacagc* |
| | ***MW70_pAaTIP1-R1*** | *cttcccacggctaccttgttcc* | ***MW72-pAaTIP1-R2*** | *gctagctcttcgtctcacattgcagcagagctcgcgcggagc* |
| ***5*** | ***AaUbi*** | | ***AaUbi-Domestication*** | |

| | | | | |
|---|---|---|---|---|
| | ***MW65_pAaUbi1-F1*** | *tagatggcaagataggccagg* | ***MW67_pAaUbi1-F2*** | *gtgagctcttcgtctctggagtgactgagcaagaattcgtgc* |
| | ***MW66_pAaUbi1-R1*** | *tggtcttcacgttgtcgatgg* | ***MW68_pAaUb i1-R2*** | *gctagctcttcgtctcacattattgcaaacttggtagacggac* |
| ***5*** | ***Genotyping Line Cam1 left*** | | | |
| | ***MW-T4_insert5p_nes-F*** | *ggttgtagacagcagtagaatcc* | ***MW-T4_insert5p_nes-R*** | *aaagtgccgataaacataacgatc* |
| | ***MW_T4_insert-F* (nested PCR)** | *gtgaagccgtaaatgacgtgatgg* | ***AA076-gen-R*** | *gagattcttcgccctccgagag* |
| ***6*** | ***Genotyping Line Cam1 right*** | | | |
| | ***MW-T4_insert3p_nes-F*** | *tgtgtgagtagttcccagataagg* | ***MW-T4_insert3p_nes-R*** | *ccagacatccatccatatttcacc* |
| | ***MW_Term_seq-F* (nested PCR)** | *gggtttcgctcatgtgttgagc* | ***MW_T4_insert-R*** | *acttgtcacctgctcactctgg* |
| ***7*** | ***Genotyping Line Cam2 left*** | | | |
| | ***MW-T8_insert5p_nes-F*** | *aaactggctcacagtacacag* | ***MW-T4_insert5p_nes-R*** | *aaagtgccgataaacataacgatc* |
| | ***MW_T8_insert-F* (nested PCR)** | *gtcacttcagctcttgctctgg* | ***AA076-gen-R*** | *gagattcttcgccctccgagag* |
| ***8*** | ***Genotyping Line Cam2 right*** | | | |
| | ***MW-T8_insert3p_nes-F*** | *gacaaccactacctgagcacc* | ***MW-T8_insert3p_nes-R*** | *aacgaatatatccactcaaatatatgcctagg* |
| | ***MW_eGFP-F2* (nested PCR)** | *cgatcacatggtcctgctgg* | ***MW_T8_insert-R*** | *gtgatgtgtagtactgctagcgc* |
| ***9*** | ***Genotyping Line BTI1 left*** | | | |
| | ***FW1prepre-F*** | *aaagcaatggtgttaacagaaagc* | ***End35S-R*** | *tgatgttggatcctctagagtcc* |
| | ***FW1pre-F* (nested PCR)** | *gcgcaggttgtgaataatgacag* | ***Mid35S-R*** | *gtagacgtggttggaacgttttc* |
| ***10*** | ***Genotyping Line BTI1 right*** | | | |
| | ***FW1prepre-R*** | *gacagcttcatcccgacgac* | ***End35S-R*** | *tgatgttggatcctctagagtcc* |

| | | | | |
|---|---|---|---|---|
| | ***FW1pre-R* (nested PCR)** | *cagtgacaatgcggaggtaga* | ***Mid35S-R*** | *gtagacgtggttggaacgttttc* |
| ***11*** | ***Genotyping Line BTI2 left*** | | | |
| | ***FW2prepre-F*** | *caaactacctggagctcccc* | ***End35S-R*** | *tgatgttggatcctctagagtcc* |
| | ***FW2pre-F* (nested PCR)** | *tcgactctccccggcaacgc* | ***Mid35S-R*** | *gtagacgtggttggaacgttttc* |
| ***12*** | ***Genotyping Line BTI2 right*** | | | |
| | ***FW2prepre-R*** | *catgtccattgtgacaaacctgc* | ***End35S-R*** | *tgatgttggatcctctagagtcc* |
| | ***FW2pre-R* (nested PCR)** | *ggatgaacgttacagtgttgtac* | ***Mid35S-R*** | *gtagacgtggttggaacgttttc* |
| ***13*** | ***Genotyping Line BTI3 left*** | | | |
| | ***FW3prepre-F*** | *gttggcttgtgccttatcttgtg* | ***End35S-R*** | *tgatgttggatcctctagagtcc* |
| | ***FW3pre-F* (nested PCR)** | *cacaccctcctaacgaaaaag ag* | ***Mid35S-R*** | *gtagacgtggttggaacgttttc* |
| ***14*** | ***Genotyping Line BTI3 right*** | | | |
| | ***FW3prepre-R*** | *taatcgtggaaaatggggcgtc* | ***End35S-R*** | *tgatgttggatcctctagagtcc* |
| | ***FW3pre-R* (nested PCR)** | *aagcctaacatgcagcagctc* | ***Mid35S-R*** | *gtagacgtggttggaacgttttc* |
